# Cryogenic Electron Tomography Redefines Herpesvirus Capsid Assembly Intermediates Inside the Cell Nucleus

**DOI:** 10.1101/2025.06.27.661840

**Authors:** Stefan L. Oliver, Muyuan Chen, Leeya Engel, Corey W. Hecksel, Xueting Zhou, Michael F. Schmid, Ann M. Arvin, Wah Chiu

## Abstract

Herpesviruses encapsulate their double-stranded DNA (dsDNA) genomes within an icosahedral nucleocapsid formed in the infected cell nucleus. Four biochemically purified nucleocapsids have been characterized, but their roles in herpesvirus replication remain controversial. The status of the capsid vertex-specific component (CVSC), essential for capsid stability and dsDNA packaging and retention, is also unclear. By integrating cryogenic focused ion beam milling with electron tomography and subtomogram averaging, we derived atomic models for all protein components, including the CVSC, across different herpesvirus capsid types within infected cell nuclei. Focused classification of pentonal vertex densities revealed differences in CVSC occupancy between genome-filled capsids and capsids lacking dsDNA, highlighting structural heterogeneity and providing insight into distinct capsid assembly stages *in situ*. These intra-nuclear findings redefine the maturation model of herpesvirus capsid assembly, advancing the understanding of herpesvirus replication, and demonstrate the effectiveness of *in situ* electron imaging by studying virus assembly within host cells.

## INTRODUCTION

The *Herpesviridae* encompass a large family of viruses comprised of alpha-, beta- and gamma subgenera that infect humans and most animal species with wide ranging health and economic consequences ^1–8^. Varicella-zoster virus (VZV), a human alphaherpesvirus, causes varicella (chicken pox) as the primary infection, and zoster (shingles) after reactivation from latency in sensory ganglia^1^, and can be life threatening in immunocompromised patients ^9–15^. VZV vaccines have reduced morbidity substantially ^16–18^ and antiviral drugs are effective but neither prevent VZV latency in sensory neurons ^19,20^ or post-herpetic neuralgia after zoster ^21,22^, and antiviral resistance can emerge ^23,24^. The other two human alphaherpesviruses, herpes simplex virus (HSV) 1 and 2 (HSV-2), also create a significant disease burden ^25,26^. HSV vaccines are not available and HSV persists in neurons despite antiviral treatment ^8^. Vulnerable populations need new interventions for these viruses and against the betaherpesvirus, human cytomegalovirus ^4^, and the Epstein-Barr and Kaposi’s sarcoma gammaherpesviruses, which cause a range of cancers ^2,3^.

Herpesviruses are classified by their distinct 4-layer morphology composed of a double stranded DNA (dsDNA) core, capsid, tegument, and envelope ^27^. These complex entities form through a convoluted morphogenesis process starting inside the infected cell nucleus. The dsDNA genome is packaged into newly synthesized capsids that exit the nucleus across nuclear membranes into the cytoplasm, where they traffic to sites of secondary envelopment, completing acquisition of tegument proteins and the lipid envelope containing virus glycoproteins. The nascent enveloped particles are transported in vesicular structures to the plasma membrane and released into the extracellular milieu. The glycoproteins protruding from the lipid membrane then mediate virus entry into neighboring cells and tegument proteins support the intracellular transport of capsids to nuclear pores where the dsDNA genome is injected; additional tegument proteins undergo nuclear translocation to initiate transcription factor functions. Since the herpesvirus capsid structure is essential for virus genome preservation during infection, high-resolution information about the progression of capsid assembly holds great potential to identify novel targets for next generation pharmaceutical interventions and vaccines.

Herpesvirus capsids are ∼125nm in diameter and share a common icosahedral symmetry with a T=16 geometry made up of 60 asymmetric units. The capsid assembles in the nucleus from 161 capsomeres, composed of 150 hexons and 11 pentons that form an icosahedron with 12 vertices, including a unique portal vertex for the packaging of the dsDNA genome and its release through the host cell nuclear pore in newly infected cells. Capsids undergo several transformative stages starting with immature pro-capsids, which are thought to form as capsomeres coalesce with the scaffold protein to produce segments and subsequently spherical capsids ^28,29^. Maturation of pro-capsids occurs as the scaffold protein is proteolytically digested by its companion protease, triggering a conformational change from a more spherical shape to their final icosahedral morphology, with 20 flat triangular faces and 30 edges. Mature capsids have been conventionally classified as A-, B-, and C-capsids, as they differ in their buoyant densities and morphologies by thin section electron microscopy (EM) ^30^. The cavities of C-capsids contain packaged dsDNA genomes, whereas B-capsids have persistent scaffold protein and its associated protease, and A-capsids are described as empty. While A- and B-capsids have been presumed to be dead-end assembly products ^29^, an alternative hypothesis is that one or both might represent assembly intermediates with the potential to encapsidate dsDNA genomes ^31^. Capsid purification can induce artefacts, such as loss of genomic DNA by disruption of the portal vertex cap, creating empty A-capsids from C-capsids ^32^. B-capsids accumulated when unit length genome formation was blocked by drug inhibition of the genome-transferase complex ^31^, suggesting that scaffold protein removal is linked to dsDNA insertion, and capsids with scaffold protein in the cavity might be capsid intermediates rather than representing a failure of genome encapsidation.

The proteins that assemble to create a mature herpesvirus capsid have orthologues in all *Herpesviridae*. These include the major capsid protein (MCP), the small capsomere interacting protein (SCP), and the capsid triplex proteins 1 (TRX1) and 2 (TRX2). The multimeric subunits formed by these proteins to produce fully mature capsids have been described at near atomic resolution for all three subfamilies of herpesviruses ^33–43^. Briefly, the MCP forms the capsomeres of hexons and pentons of the icosahedral capsid. Six copies of the SCP decorate the tops of the hexons. Heterotrimeric triplexes (TRX1 monomer and TRX2 dimer) occupy the canyons between the capsomere towers and cover up the holes in the MCP floor to stabilize the capsid for dsDNA genome containment ^33,44^. At each of the 12 capsid vertices, five copies of the capsid vertex specific component (CVSC) bind to triplexes adjacent to the pentons and the unique portal vertex ^33^. In HSV, the CVSC is a heteropentamer composed of orthologues shared with VZV; the capsid vertex-specific component 1 (CVC1) monomer, two copies of CVC2, and two copies of the protein designated VP1/2. CVSC binding has been associated with reinforcing capsid stability to withstand the very high packaging pressure of the dsDNA genome ^45–47^. At the portal vertex, a dodecamer (VZV ORF54; HSV-1 pUL6), replaces a penton with a portal and tentacle-like helices that ring the DNA translocation channel, and an ATP-driven terminase is recruited to cleave the herpesvirus dsDNA genome for packaging ^29,34–36,38,45,48–50^. The CVSC is expected to form part of the portal cap that retains the dsDNA genome in the capsid ^29,34–36,38,45,48–50^. The CVSC heteropentamer has also been called the capsid-associated tegument complex because its subunits include the large tegument protein (VP1/2), VZV ORF22 and HSV pUL36 ^51^; in HSV and PRV, these orthologues have been identified as inner tegument proteins that function in microtubule-mediated capsid transport to cell nuclei ^52,53^. The CVSC of CMV, a betaherpesvirus, has a non-conserved tegument subunit, pp150 ^42,54^, while gammaherpesvirus CVSCs are thought to resemble those of the alphaherpesviruses ^51^.

Critical information about the structural composition of herpesvirus capsids was gained using cryogenic-electron tomography (cryo-ET) of purified particles but resolution was limited ^55–63^. The adaptation of single particle cryo-EM enabled capsid structures of most human herpesviruses to be determined at better than 4Å resolution ^33,35–41,47^. However, knowledge about capsid assembly as it occurs *in situ* within the infected cell nucleus remains limited. The CVSC has been proposed to associate with the capsid inside the infected cell nucleus, but the structural organizations of their individual components have not been visualized directly in the nucleus of herpesvirus infected cells ^64,65^. Whether the assembly of the VP1/2 subunit of the CVSC takes place in the nucleus or cytoplasm is unclear ^64,66,67^. Furthermore, despite previous high-resolution structures for VZV capsids, the CVSC and the portal vertex were not resolved in these studies ^39,41^. Recent work with purified capsids partially resolved the CVSC and the portal vertex ^68^. To better understand capsid assembly and the relationship between capsids conventionally classified as A- and B-capsid types compared to the dsDNA containing C-capsids, we sought subnanometer resolution imaging of capsids within VZV infected cells. Imaging inside mammalian cells is challenging because cell thickness limits the penetration of the electron beam and reduces the image contrast due to inelastic scattering and multiple electron scattering ^69^. Even at high voltages (1MV), the electron beam can only penetrate 1µm; the nuclei of typical mammalian cells range in size from 5 to 10µm. To overcome this limitation, recent advances in cryogenic focused ion beam and scanning electron microscopy (cryo-FIB/SEM) have been introduced to mill away most of the thick and vitrified cell, leaving a thin lamella of 100 to 400 nm in thickness. When integrated with cryo-ET using a 300 keV electron microscope, 3D volumes can be produced at near-atomic resolution by imaging at different angles, cryo-ET reconstruction, and subtomogram averaging (STA) ^70^. We applied this approach to visualize VZV particles within infected cells in a frozen, hydrated state, without the use of contrast agents, and determined the structures of their component proteins.

Our findings reveal that the CVSC heteropentamer assembles on capsids in the nucleus of infected cells and that the definition of the asymmetric unit of the icosahedral VZV capsid should be expanded to incorporate the CVSC interactions with neighboring MCPs and triplexes, which were absent from previous capsid models ^39,41^. Focused classification, including of the unique portal vertex, indicates that genome-filled C-capsids have full CVSC occupancy, defined as the maximum of five heteropentamers. Among capsids lacking dsDNA, empty A-capsids were not detected within the infected cell nucleus whereas capsid assembly intermediates (CAIs) could be distinguished by differences in the number of CVSCs found at the vertices of the icosahedron, which correlated with levels of scaffold protein in the capsid cavity, suggesting a continuum along the assembly pathway. Notably, CAIs would be grouped together as B-capsids based on scaffold protein alone. These observations provide evidence that CVSC acquisition can serve as a temporal marker for the progression of intra-nuclear herpesvirus capsid assembly *in situ*, consistent with its biological importance for ensuring the stability of genome-containing capsids during nuclear egress and formation of complete virus particles in the cytoplasm.

## RESULTS

### VZV morphogenesis visualized within infected cells *in situ* by cryo-FIB-SEM and cryo-ET

To visualize herpesvirus morphogenesis in 3D-space, a workflow was developed to distribute VZV infected cells evenly on EM grids suitable for cryo-FIB milling. First, because VZV does not yield high titer cell-free virus particles ^1^, it was critical to optimize the passage of infected cells onto uninfected cells to initiate infection prior to vitrification. Infection was monitored by the fluorescence RFP signal produced by MeWo cells infected with a recombinant VZV (pOka-TK-RFP ^71^) expressing a thymidine kinase (TK) tagged with a monomeric red fluorescence protein (RFP; **Fig. 1**); the pOka-TK-RFP maintains the characteristics of the bacterial artificial chromosome derived WT pOka ^71–74^. Second, to overcome the propensity of VZV infected cells to adhere to the grid bars when seeded on SiO_2_ EM grids, islands of extracellular matrix were micropatterned within the grid squares (**Fig. 1b**). With this technique, infected cells were positioned centrally within the grid squares after vitrification and ideal for cryo-FIB milling to produce lamella suitable for cryo-ET (**Fig. 1c and 1d**). Tomograms of cryo-FIB milled cells revealed the complex cellular environment in VZV infected cells (**Fig. 1e; Suppl. Fig. 1 to 7; Suppl. Table 1 and 2**). Cellular features were readily visible in medium magnification (16.9 to 39.1Å/pixel) montage maps used to facilitate cryo-ET tilt series data collection from cells thinned to a target thickness of ∼175nm; the nuclear envelope with visible nuclear pores demarcated the intranuclear and cytoplasmic compartments (**Fig. 1d; Suppl. Fig. 1a, 3a to 7a**). A plethora of vesicular structures, mitochondria, and, within infected cells, VZV capsids and complete virus particles, was visible, facilitating the localization of regions of interest in a tilt series collection. The higher resolution tomographic reconstructions yielded substantial contrast enabling machine learning-based segmentation to identify ribosomes, microtubules, microfilaments, complex vesicular structures, and assembled capsids in the nucleus and sites of virus particle morphogenesis in the cytoplasm (**Suppl. Fig. 7c**).

**Figure 1.**
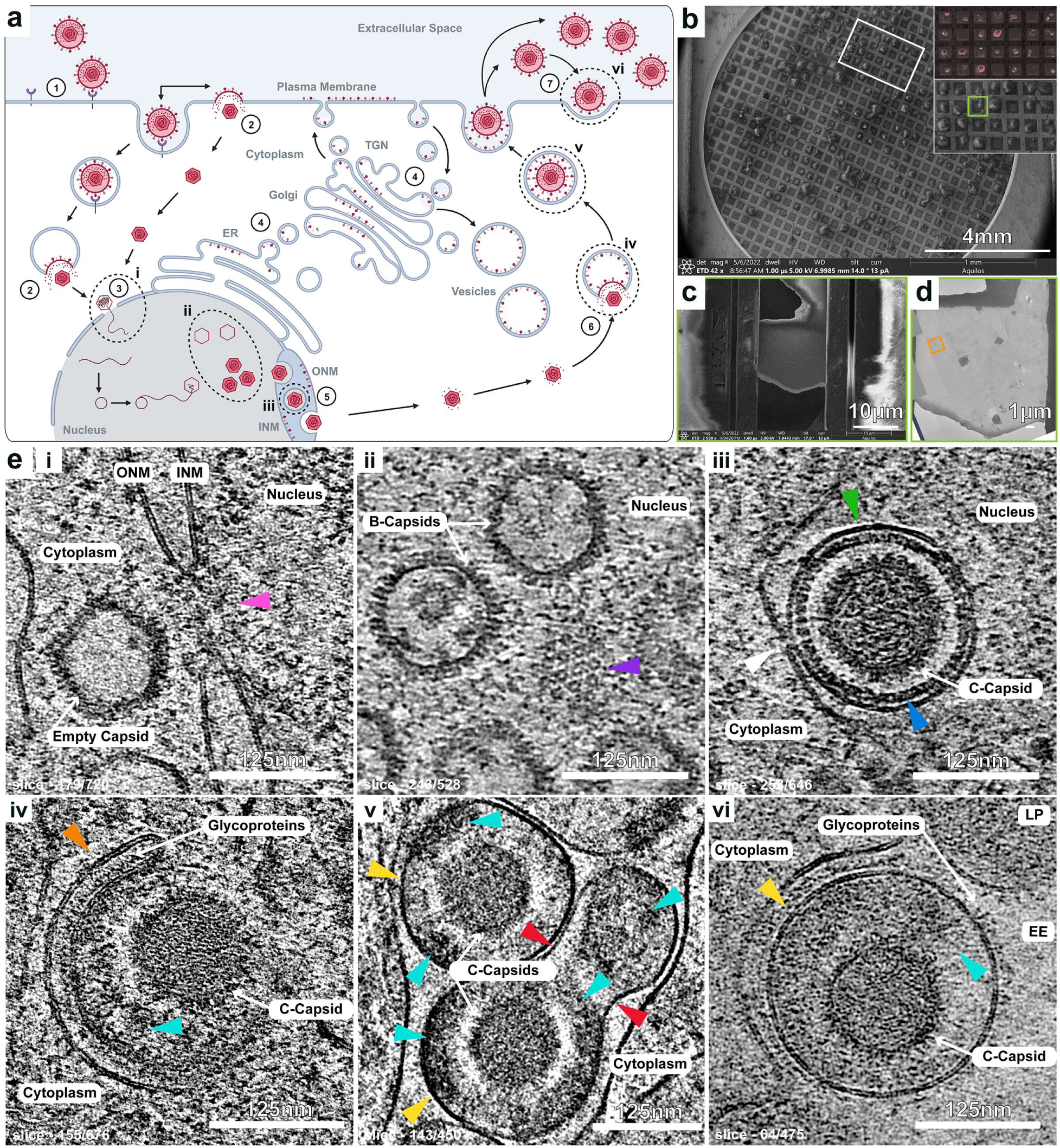
The VZV replication cycle and stages of morphogenesis within infected MeWo cells in 3D space captured by cryo-FIB/cryo-ET. **(a)** A cartoon representation (drawn using Biorender) of the VZV replication cycle where the key steps are numbered from 1 through 7 (black circles). VZV glycoproteins bind cell surface receptors during virion attachment (1), allowing the gB/gH-gL fusion complex to fuse the envelope with cell membranes either in endocytic vesicles or directly with the plasma membrane (2). The capsid is released into the cytoplasm, traffics to and docks with a nuclear pore where the dsDNA genome is injected into the cell nucleus (3) to initiate replication and capsid assembly. VZV glycoproteins necessary for attachment and cell entry, are synthesized in the endoplasmic reticulum (ER) and trafficked to the Golgi during maturation, exocytosed then endocytosed and trafficked to the trans-Golgi network (TGN) or trafficked to the nuclear envelope (inner nuclear membrane, INM; outer nuclear membrane, ONM) (4). Newly synthesized capsids undergo nuclear egress through the nuclear envelope via envelopment and de-envelopment courtesy of the nuclear egress complex (5). Glycoproteins are trafficked from the TGN to sites of secondary envelopment, where capsids acquire additional tegument proteins, and incorporated into nascent virus particles (6). VZV particles are exported in vesicular structures to the extracellular environment (7) where they can either be released into the extracellular environment or attach to the plasma membrane. **(b)** Cryogenic scanning electron microscopy (cryo-SEM) of VZV infected MeWo cells seeded on a micropatterned Quantifoil 300 mesh SiO_2_ grid. The inset shows a zoomed in image of fluorescence microscopy (right top) of the grid prior to vitrification (red fluorescence indicates infection with recombinant VZV expressing TK-RFP) and the cryo-SEM image post vitrification (right bottom). The green square highlights a single cell where a lamella was milled and presented in panels **c** and **d**. **(c)** A lamella produced by cryo-FIB milling of the VZV infected cell highlighted in A. **(d)** A montage image (16.85Å/pixel) produced by cryogenic transmission electron microscopy (cryo-TEM) of the lamella from panel **C**. The inset orange box highlights the zoomed in area where tilt series data were collected to generate the tomogram in panel Ev. **(e)** In our cryo-FIB/cryo-ET data, we have captured capsids corresponding to some of the VZV replication stages labeled in Roman numbers (i-vi) and outlined by dotted black ellipses in panel A. Each image is a single slice through a low pass filtered tomogram. **(i)** An empty VZV capsid is visible at the nuclear pore where the inner and outer nuclear membrane converge (pink arrowhead). **(ii)** B-capsids containing scaffold protein and the capsomeres of an adjacent capsid along the 3-fold axis (blue violet arrowhead). **(iii)** A C-capsid surrounded by the nuclear egress complex (navy blue arrowhead) and the primary envelope (lime green arrowhead) undergoing fusion with the outer nuclear membrane (white arrowhead). **(iv)** A C-capsid interacting with the tegument layer (bright turquoise arrowhead) and glycoprotein-coated cell membranes (tangerine arrowhead) morphing around the capsid-tegument complex. **(v)** VZV particles containing C-capsids, one that has undergone complete morphogenesis and a second in the process of the final steps of morphogenesis and the production of a light particle. The tegument is clearly visible as the dark structures (bright turquoise arrowheads) between the capsids and the VZV particle envelope (gorse yellow arrowheads). Red arrowheads indicate a point of constriction between the formation of a complete VZV particle and a light particle. **(vi)** VZV particles released into the extracellular environment (EE) bound to the plasma membrane. One contains a C-capsid with the tegument (bright turquoise arrowheads) clearly visible as the dark structures between the capsids and the VZV particle envelope (gorse yellow arrowheads), which is coated with glycoproteins. The second is a partially visible light particle (LP) also bound to the plasma membrane. Scale bars 125nm.

Several critical steps in the herpesvirus replication cycle were captured within cryo-ET tomograms generated from cryo-FIB milled cells infected with VZV (**Fig. 1e; Suppl. Fig. 1 to 8; Suppl. Movie 1 to 6**). These included capsids docked at the nuclear pore (**Fig 1ei; Suppl. Fig 1; Suppl. Movie 1**), newly synthesized capsids in the nucleus (**Fig 1eii; Suppl. Fig 3; Suppl. Movie 2**), nuclear egress (**Fig 1eiii; Suppl. Fig 4; Suppl. Movie 3**), secondary envelopment (**Fig 1eiv; Suppl. Fig 5; Suppl. Movie 4**), VZV particle morphogenesis (**Fig 1ev; Suppl. Fig. 6 and 7; Suppl. Movie 5**), and VZV particles bound to the extracellular plasma membrane surface (**Fig 1evi; Suppl. Fig 8; Suppl. Movie 6**), a typical feature of this cell associated virus ^1^. Within the infected cell nucleus, pro-capsids and A-, B-, and C-capsids based on conventional morphological features, were distinguishable in 3D-space at this resolution. Procapsids were spherical and contained a solid density of scaffold protein; capsids apparently devoid of scaffold protein or dsDNA (conventional A-capsids) were identified; B-capsids were observed and defined by the presence of scaffold protein (**Fig. 1eii**); and C-capsids were distinguishable by their cavities filled with dsDNA (**Fig. 1eiii, 2A and 2B; Suppl. Fig. 4 to 6**). Procapsid assembly precursors were identified within infected cell nuclei, appearing as angular segments of capsomeres associated with scaffold protein with a curvature similar to the procapsids (**Suppl. Fig. 2**). Importantly, high-resolution detail was evident by the visualization of individual capsid capsomeres (**Fig 1eii, Suppl. Fig 3; Suppl. Movie 2**), individual nuclear egress complexes (**Fig 1eiii, Suppl. Fig 4; Suppl. Movie 3**), and individual glycoproteins on enveloped VZV particles in the cytoplasm or bound to the plasma membrane (**Fig. 1eiv and vi; Suppl. Fig. 5, 8 and 9; Suppl. Movie 5 and 6**). Partially assembled virus particles were observed within cytoplasmic sites of VZV secondary envelopment and morphogenesis, including sites of tegument protein accumulation, based on the electron dense protein surrounding the capsids within the reconstructed tomograms (**Fig. 1eiv and 1ev; Suppl. Fig. 5 and 6**). For example, a VZV light particle, a glycoprotein-studded envelope containing tegument proteins but without a capsid, and a complete particle were observed in a vesicular structure, and in a neighboring vesicular structure, an elongated VZV particle contained a C-capsid and the accumulation of tegument protein associated with a membranous structure (**Suppl. Fig. 7b**). The virus envelope within these vesicular structures was detected by machine learning-based segmentation (**Suppl. Fig. 7c**). Collectively, these direct visualizations reveal details of the VZV replication cycle previously unobserved in its native hydrated state inside infected cells, particularly for nuclear egress, tegument acquisition during secondary envelopment, and virus glycoprotein interactions with both intra- and extracellular membranes.

### Subnanometer reconstruction of VZV capsid proteins in infected cells

From 194 tilt series, 1,084 capsids were identified and classified based on classical morphological features, the presence or absence of scaffold protein or the virus genome in the capsid cavity. A visual inspection of the image contrast in the capsid cavity revealed a continuum of sparse density, likely scaffolding, suggesting that completely empty A-capsids are absent or very rare within infected cell nuclei (**Suppl. Fig. 10**). Residual scaffold protein was limited enough in some capsids to be obscured by conventional EM techniques due to the plane of the cut in ultrathin sections in previous studies. The cellular distribution of the 1,084 VZV capsids was primarily localized inside the nucleus (n=980) with only 9.3% detected in the cytoplasm either as naked capsids, without tegument or envelope proteins (n=42), or as complete enveloped VZV particles (n=59) (**Suppl. Table 3**). The capsid distribution was typical for peak particle synthesis of VZV infected cells, which aligns with thymidine kinase (TK) expression, an early gene, as determined by the red fluorescence signal produced from the recombinant pOka-TK-RFP virus.

An initial 13Å cryo-ET map of the VZV capsid was generated by subtomogram averaging (STA) of 317 capsids (**Fig. 2; Suppl. Fig. 11**), containing variable quantities of scaffold protein or packaged DNA. This icosahedral symmetry-imposed map clearly shows the T=16 icosahedron, composed of 60 asymmetric units. Each asymmetric unit consists of a single pentonal MCP subunit (P^P^), one peripentonal hexon (P), one central hexon (C), and half of an edge hexon (E), along with interdigitating triplexes (Ta, Tb, Tc, Td, and Te) (**Fig. 2c and 2d**). Cryo-ET maps were produced for the vertex around the 5-fold axis (8.3Å) and the 3-fold symmetry axis (8.4Å) by subsequent symmetry expansion and focused refinement of the 317 capsids. (**Fig. 2e and 2f; Suppl. Fig. 11**). The 8.3Å vertex map had sufficient resolution to perform segmentation of individual proteins for the pentonal MCP (P^P^), six MCPs (P^1–6^) for the peripentonal hexon and the associated small capsomere interacting protein (SCP), triplex proteins TRX1 (monomer) and TRX2 (dimer) that form the α_2_β heterotrimers of Ta and Tc, and the three proteins, CVC1 (ORF43 monomer), CVC2 (ORF34 dimer), and ORF22 dimer, that assemble to form the capsid vertex specific component (CVSC) heteropentamer (**Fig. 3 and 4; Suppl. Table 4**). Similar to the vertex map, individual proteins were segmented for the 8.4Å 3-fold symmetry axis cryo-ET map for the six MCPs (C^1–6^) for the central hexon and associated SCP in addition to the TRX1 and TRX2 proteins of the Te triplex (**Suppl. Fig. 12**). The densities in both 3- and 5-fold axes of symmetry were segmented using the structure derived from single particle cryo-EM of VZV capsids (7BW6)^39^. All of the features for the pentonal, peripentonal hexon, and central hexon MCPs were apparent, including the tower, consisting of the crown and buttress regions (**Fig. 3a to c; Suppl. Fig. 12**), and the HK97 phage capsid protein-like fold facing the interior chamber of the capsid (**Fig. 3d**), which is evolutionary conserved for herpesviruses and dsDNA bacteriophages ^75^. The resolution of these regions in the cryo-ET map was supported by the ability to observe densities for alpha helices in each of the MCP, TRX1 and TRX2 proteins and corresponding Map-Q scores (**Fig. 3c; Suppl. Fig. 12c and e; Suppl. Fig. 13; Suppl. Movie 7**). The classical HK97 phage capsid protein fold of the MCP was distinguishable with the long alpha helix (helix 4) interdigitating at the pseudo 3-fold axis of symmetry between the penton and neighboring peripentonal hexon MCPs (**Fig. 3d**). These subnanometer resolution structural details of the MCP, SCP, TRX1 and TRX2 are the first to be visualized inside the nucleus of a herpesvirus-infected cell.

**Figure 2.**
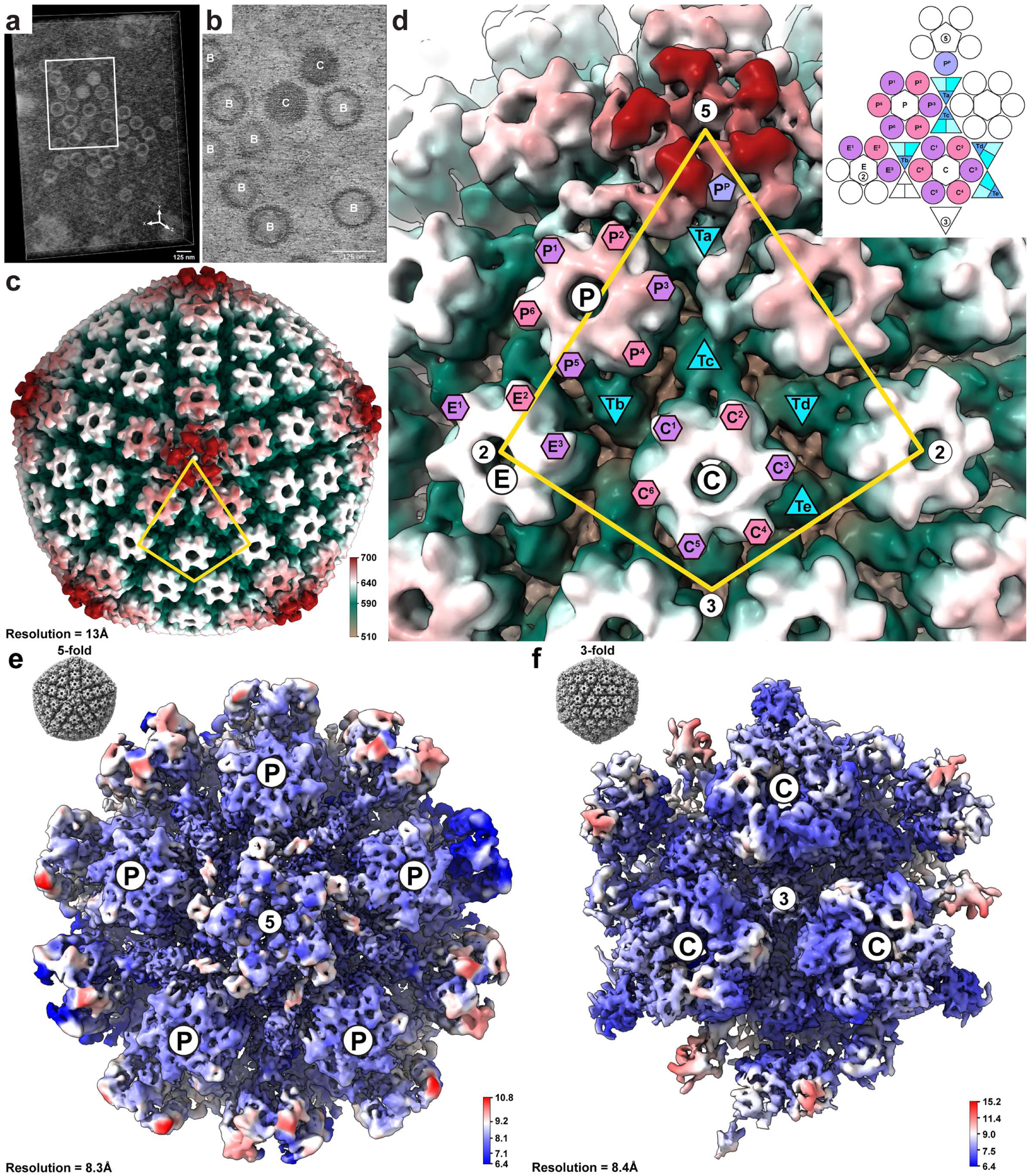
Cryo-ET density maps of the VZV capsid, capsid vertex, and the 3-fold symmetry axis. **(a)** A representative tomogram reconstruction produced from cryo-ET of a cryo-FIB/SEM milled region inside the nucleus of a vitrified, VZV infected MeWo cell represented as a 3D-volume. **(b)** A single slice from the tomogram in (**a**) in a region inside the nucleus of a vitrified VZV infected MeWo cell where a cluster of B-capsids (scaffold protein and without dsDNA labeled as B) and C-capsids (dsDNA genome labeled as C) were identified. Scale bars indicate 125nm. **(c)** Surface representation of a cryo-ET map (13Å) generated by subtomogram averaging with imposed icosahedral symmetry of 317 VZV capsids with a mixture of B and C capsids. The capsid is colored radially (510-700Å form the center of the capsid). The yellow kite connects the 2-, 3-, and 5-fold axes of symmetry. **(d)** Zoom in view of a portion of (**c**) annotated with protein components that make up an asymmetric unit composed of the pentonal major capsid protein (P^P^), the three types of hexon (peripentonal (P1-P6), edge (E1-E3), and center (C1-C6)), and the five triplexes (Ta, Tb, Tc, Td, and Te). The inset (right) is a schematic of the asymmetric unit of the icosahedral capsid. **(e and f)** Cryo-ET maps generated by subtomogram averaging and focused classification of the VZV capsid vertex (e; 8.3Å; EMD-49591) and 3-fold axis of symmetry (f; 8.4Å; EMD-49886). The inset cryo-ET maps of the 13Å VZV capsid cryo-ET map are projected along the 5-fold (**e**) and 3-fold (**f**) axes of symmetry. Peripentonal (P) and center (C) hexons are labelled. The color key indicates the local resolution (Å) of the cryo-ET maps.

**Figure 3.**
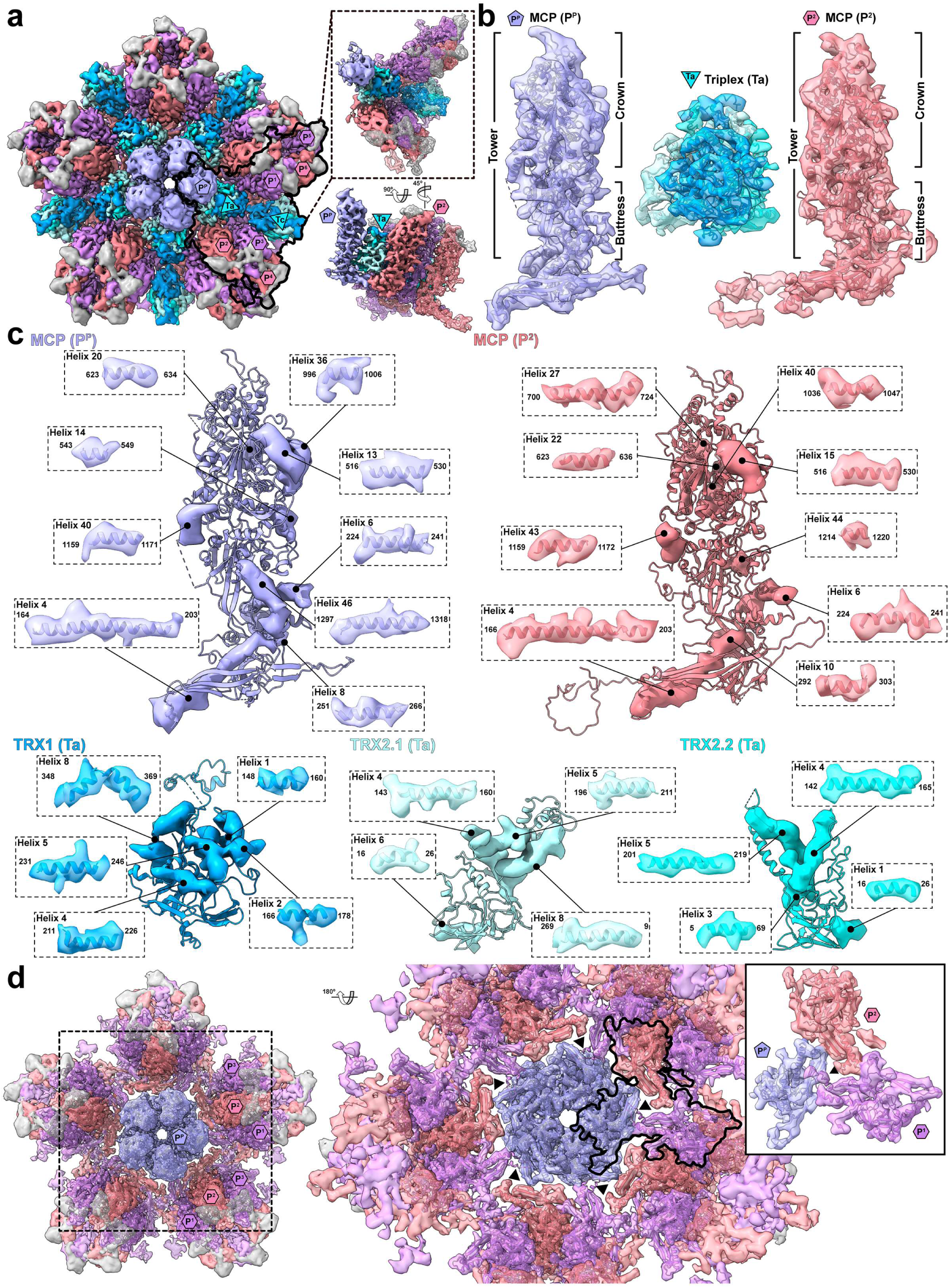
Segmentation of the 8.3Å VZV capsid vertex cryo-ET map. **(a)** Segmentation of the 8.3Å cryo-ET map based on the 7BW6^39^ and a refined model based on the subnanometer map. The major capsid protein (MCP; ORF40) of the penton (P^P^) and the peripentonal hexons (P^1–6^), triplex proteins TRX1 (ORF20) and TRX2 (ORF41) for triplexes Ta and Tc, and small capsomere interacting protein (SCP; ORF23) are colored lavender blue (P^P^), very soft violet (P^1,3,5^), light coral (P^2,4,6^), deep sky blue (TRX1), Columbia blue (TRX2.1), aqua (TRX2.2), and dark gray (SCP). The colored shapes within the black outline demarcate a portion of the asymmetric unit (P^P^, P^1–6^ and associated SCP, and triplexes Ta and Tc) of the VZV capsid vertex, which is isolated in the dotted box (inset). **(b)** Extracted densities of the P^P^ and P^2^ MCP with the tower, buttress and crown highlighted, and Ta triplex. **(c)** Exploded views of extracted densities for helices from the P^P^ and P^2^ MCP, and Ta triplex. Each helix is numbered according to the primary structure along with the beginning and end numbered amino acids. **(d)** Segmentation of the VZV capsid vertex without the triplexes. P^P^ and P^1–3^ MCPs are identified with colored shapes. The dashed boxed outlines the region shown in the right hand panel. Triangles represent the local 3-fold symmetry axes where the pentonal MCP (P^P^) intersects the two peripentonal hexon MCPs, P^1^ and P^2^. The black outline highlights the intersection between the HK97-like fold and the P^P^, P^1^, and P^2^ MCPs (inset).

**Figure 4.**
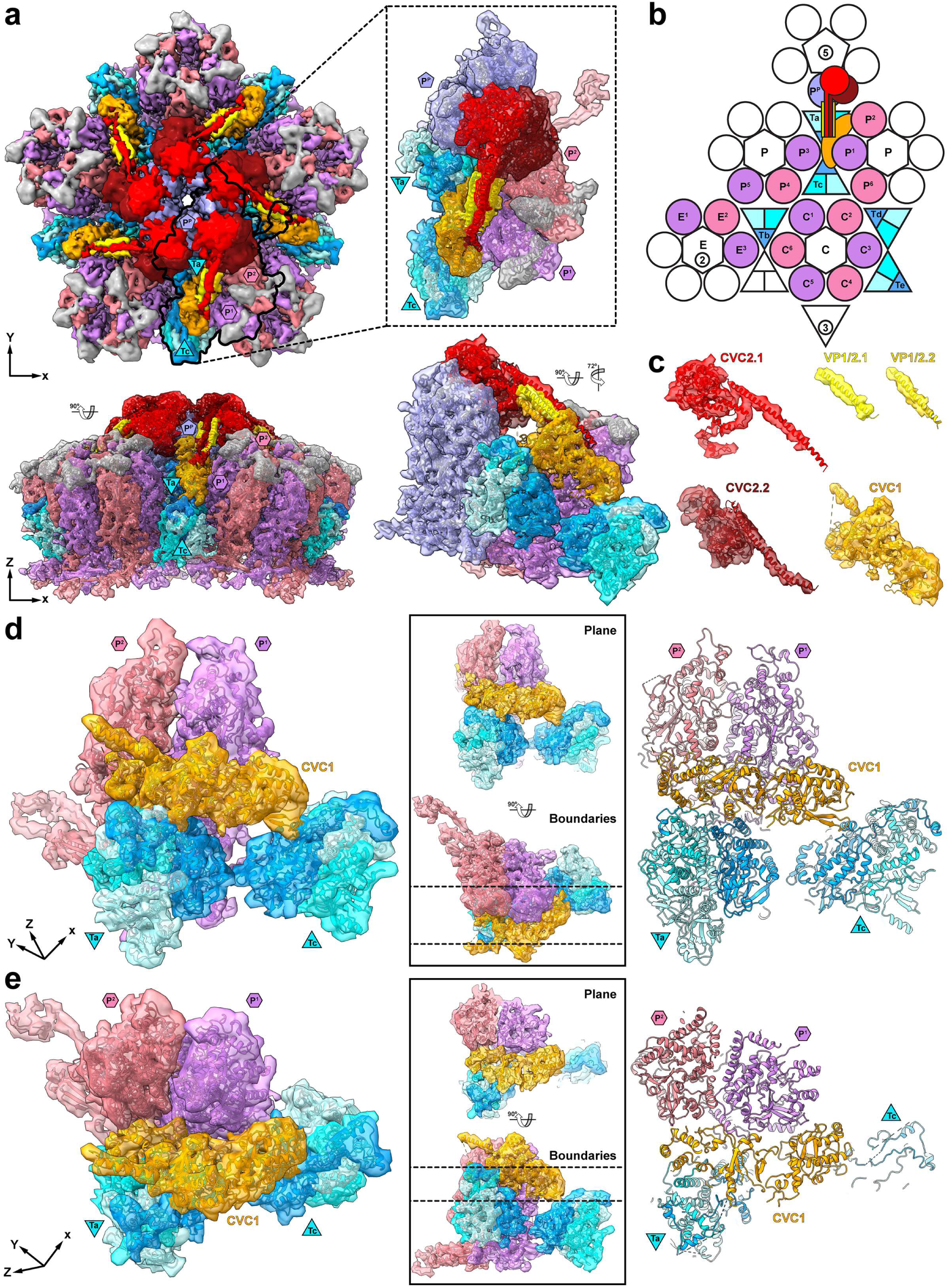
The VZV capsid vertex specific component redefines the asymmetric unit. **(a)** Segmentation of the 8.3Å cryo-ET map of the vertex around the 5-fold axis of the capsid to isolate the CVSC and fit an AlphaFold 2 model using MDFF simulation and the 7BW6^39^ model. The MCP, SCP, TRX1, and TRX2 are colored as for Fig. 3 and the CVSC subunits colored orange (CVC1; ORF43), red (CVC2.1; ORF34), falu red (CVC2.2; ORF34), yellow (VP1/2.1; ORF22), and gold (VP1/2.2; ORF22). The black outline highlights a portion of the VZV capsid vertex with the CVSC, which is isolated in the dashed box (inset). The lower panels show side views of the complete vertex map and the extracted region in the dotted box. **(b)** A cartoon representing the CVSC components’ locations in relation to the redefined asymmetric unit compared to Fig 2C. Colored shapes define the pentonal major capsid protein (P^P^), the three types of hexon (peripentonal (P^1^-P^6^), edge (E^1^-E^3^), and center (C^1^-C^6^)), the five triplexes (Ta, Tb, Tc, Td, and Te), and the CVSC. **(c)** Exploded view of the CVSC subunits. **(d and e)** Extracted densities revealing (**d**) CVC1 (ORF43) interactions with triplexes Ta and Tc, and (**e**) peripentonal hexon MCP P^1^ and P^2^. Models of the CVC1, TRX1, TRX2, P^1^, and P^2^ within the cryo-ET map densities are shown in the left-hand panels. Right hand panels show plane views of the molecular models (far right) that highlight likely interaction sites between CVC1, the triplexes (**d**), and peripentonal MCPs (**e**) and the boundaries of the planes (black boxes).

### The CVSC heteropentamer assembles as part of the capsid asymmetric unit in infected cell nuclei

The CVSC status as a capsid component has been difficult to assess due to its propensity to disassociate from vertices during herpesvirus capsid purification, including VZV ^45,76,39,41^. In contrast, the CVSC density was apparent at all 12 vertices of the capsid structure in infected cell nuclei (**Fig. 2c**). Furthermore, the 8.3Å cryo-ET map of the VZV capsid vertex was of sufficient resolution to extract densities for the CVSC, demonstrating that the heteropentamer is a capsid component acquired independently of the full tegument layer of the virus particle (**Figs. 2e, 4a and 4c**). An AlphaFold model of the CVSC structure oriented the head group for each CVC2 (ORF34) molecule at a downward angle compared to the extracted CVSC cryo-ET map density (**Suppl. Fig. 14a and 14b**). The CVC2 head group AlphaFold models were fitted into the cryo-EM density with the aid of molecular dynamics flexible fitting (**Suppl. Fig. 14c; Suppl. Movie 8**), enabling segmentation of the cryo-EM density into the five individual subunits of the CVSC heteropentamer (**Fig. 4a to 4c**). This revealed that the CVC1 subunit (ORF43 monomer), which has a limited interaction with TRX1 from triplex Te, primarily associates with all three Ta triplex subunits along with MCP P^1^ and P^2^ of the peripentonal hexon from the neighboring asymmetric unit (**Fig. 4b, 4d and 4e**). A helical bundle forms between the CVC1 subunit, the two ORF22 proteins and the two CVC2 proteins, enabling the CVC2 head group to interact with the peripentonal hexon facing side of the pentonal MCP crown (**Fig. 4a**). Since almost all capsids analyzed were intra-nuclear, these observations definitively demonstrate that CVSC assembly at the capsid vertices occurs inside the nucleus. Furthermore, these findings enabled us to redefine the asymmetric unit of the mature C-capsid to be composed of six C hexon subunits (MCP and SCP), three E hexon subunits, and a pair of three peripentonal hexon subunits from neighboring hexons, one pentonal MCP, five interdigitating triplexes, and a single CVSC heteropentamer (**Fig. 2d and 4b; Suppl. Fig. 15a**). This arrangement differs from previous models of alphaherpesvirus asymmetric units, where the interaction between CVC1 with triplexes Ta and Tc and MCP P^1^ and P^2^ were not used to define the asymmetric unit, proteins were missing, or proteins from a neighboring asymmetric unit were included (**Suppl. Fig. 15**).

### CVSC acquisition redefines capsid assembly intermediates formed during herpesvirus capsid assembly

Given uncertainties about the stages of capsid assembly based on studies of purified capsids ^45,76,77^ ^31,78^, we investigated the distribution of CVSC heteropentamers at vertices of capsids in infected cells to determine whether the CVSC occupancy could be used to differentiate capsid types and achieve a more precise characterization for the stages of capsid assembly (**Fig. 5**). Uniform Manifold Approximation and Project (UMAP) is a dimension reduction technique that can be used for the visualization and categorization of complex data ^79^, including structural heterogeneity in single particle Cryo-EM ^80,81^. Application of this unsupervised computational approach (**Fig. 5a**) using three-axis orthogonal projection in latent space classified the 317 capsids that were used to generate the 8.3Å vertex cryo-ET map (**Fig. 2**). Each capsid image was aligned to its symmetry axes, and icosahedral symmetry was applied to individual particles before the projections were generated to compensate for the missing wedge effect intrinsic to cryo-ET data ^82^. Three clusters were differentiated in latent space, revealing limited versus (i) extensive (ii) scaffold protein in the capsid cavity, and those with packaged DNA (iii) corresponding to C-capsids, as viewed along the 2-, 3-, and 5-fold axes of symmetry and the STA for each class (**Fig. 5a to 5c**). The detection of limited quantities of scaffold protein for class i capsids enabled the differentiation from conventionally designated B-capsids. The absence of any density inside the capsid cavity, previously established for classical empty A-capsids based on conventional negative stained images ^30^, was not detected within infected cell nuclei. Residual scaffold protein was visible in the native structures preserved by vitrification performed in this study (**Suppl. Fig. 10**). In addition to capsid cavity content differences, the three UMAP capsid clusters were found to differ in CVSC occupancy at the vertices (**Fig. 5d to 5f**). For each of the 317 capsids, sub-particles were extracted from the 12 vertices and aligned to the same C5 symmetry axis. These particles were initially subject to binary reference free focused classification, resulting in two classes reflecting vertices without or with CVSC occupancy (**Fig. 5d and 5e**). The CVSC was either absent or present, with the arms and lobes of the CVSC clearly visible in the C5 symmetry STA maps (**Fig. 5d**). Capsid vertices were then classified into those without and with the CVSC and by their UMAP cluster, i to iii (**Fig. 5e**). As expected, CVSC occupancy was highest on C-capsids in infected cells, as has been reported for purified capsids ^33,37,40,42,47^, and because they are considered complete and ready for egress from the nucleus for final tegumentation and envelopment in the cytoplasm ^45^. Persistent scaffold protein is the conventional marker for B-capsids by EM, resulting in greater buoyant density than empty A-capsids ^30^. Notably, capsids differed in CVSC occupancy depending on whether scaffold protein was limited or extensive within the capsid cavity, with the former having greater CVSC occupancy than the latter (**Fig. 5b to 5f**). Critically, for the 12 vertices of individual capsids, CVSC occupancy at each vertex increased with decreasing levels of scaffold protein density (**Fig. 5f**). Since C-capsids have the highest CVSC occupancy, these findings indicate that capsids with relatively more CVSC occupancy and less scaffold protein are capsid assembly intermediates (CAIs) further along the pathway of capsid maturation than those with less CVSC occupancy and more residual scaffold protein.

**Figure 5.**
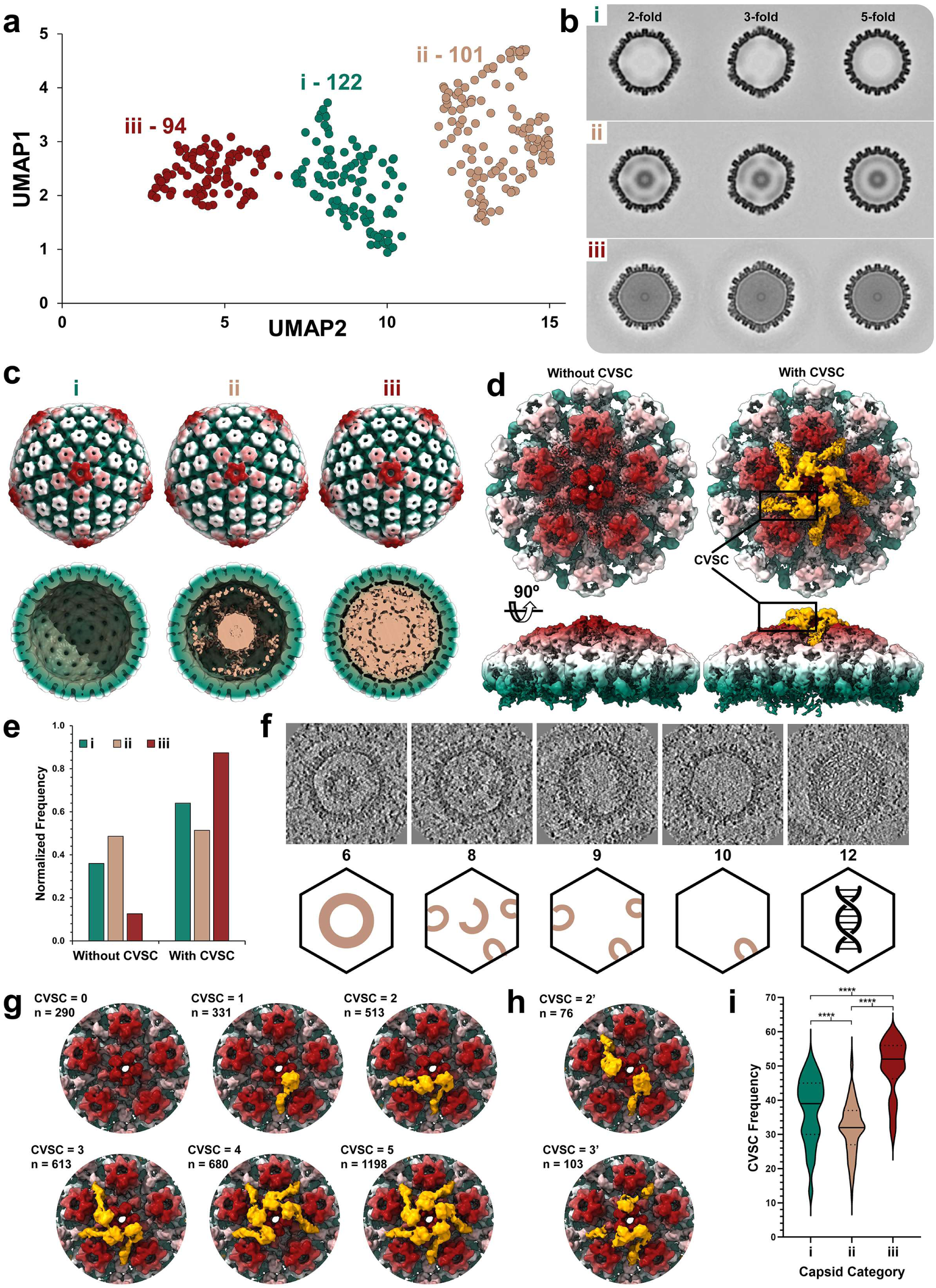
The capsid vertex specific component as a temporal marker for herpesvirus capsid assembly. **(a)** Uniform manifold approximation and projection (UMAP) of 317 VZV capsids producing three clusters, i, ii and iii. **(b)** 2D class averages of clusters i to iii projected along 2-, 3-, and 5-fold axes of symmetry. **(c)** Subtomogram reconstruction of the three capsid classes, i, ii, and iii. Capsids are colored radially from the center of the capsid to the outer shell; beige, green, white, and red. The transverse planes show the empty cavity, (i), scaffold filled cavity (ii), and DNA filled cavity (iii) of the VZV capsids. **(d)** Subtomogram reconstructions of the capsid vertex without and with the CVSC. The vertices are colored radially as for panel B with the CVSC colored yellow. **(e)** Normalized frequency of i, ii, and iii capsid classes without or with the CVSC. **(f)** Increased CVSC occupancy at capsid vertices associated with decreased scaffold protein. Slice view of representative polished particles (EMAN2) extracted from tomograms and cartoons depicting the scaffold protein visible in the capsid cavity. Numbers represent the frequency of vertices occupied with the CVSC. **(g)** Subtomogram averaging after symmetry relaxation and focused classification. The frequency (n) of VZV capsid vertices without (0) or with 1 to 5 copies of the CVSC (yellow) is indicated for each vertex. **(h)** Non-consecutive distribution of the CVSC (yellow) at the VZV capsid vertices with the frequency (n) of 2 (2’) or 3 (3’) copies after subtomogram averaging and focused classification. **(i)** A violin plot representing the frequency of the CVSC (60 copies per capsid; 5 CVSC subunits per 12 vertices of the capsid) for each capsid class, i, ii and iii. The solid line represents the median and the dotted lines represent quartiles. Asterisks above the violin plots indicate p-values (<0.0001) for significant differences from a Tukey’s multiple comparison test.

To further scrutinize the CVSC on capsids in infected cells, vertex particles were subjected to C5 symmetry expansion, generating five sub-particles for the asymmetric orientations. Additional focused classification using a mask covering a single CVSC site produced two classes of CVSC occupancy at each individual location, those with or without the heteropentamer. This procedure established the frequency of CVSC heteropentamers, producing vertex sub-particles with either zero copies, 1 to 5 copies, or 2 and 3 copies of non-consecutive CVSC distribution, which were used for STA without symmetry imposed (**Fig. 5g and 5h**). Each capsid has a potential maximum of 60 copies of the CVSC heteropentamer, as there are 12 capsid vertices and 5 CVSC per vertex, including the portal vertex ^29,34–36,38,45,48–50^. For the 317 capsids in the present study, there were 3,804 vertices. Frequencies ranged from 290 vertices without a CVSC to 1,198 vertices with a full complement of five copies of the CVSC (**Fig. 5g**). Additionally, alternative distributions of the CVSC were observed at vertices where 2 or 3 copies of the heteropentamer were not adjacent to one another, indicating a non-cooperative attachment of the heteropentamer to these capsid vertices (**Fig. 5h**). In contrast to purified VZV C-capsids ^41^, intranuclear C-capsids in infected cells had almost complete vertex occupancy (52; SEM = 0.861), demonstrating their maturity (**Fig. 5i**). CAIs with limited scaffold protein had greater CVSC occupancy (39; SEM = 0.859) compared to CAIs with extensive scaffold protein (32; SEM = 0.753), providing additional evidence that the former represent a stage closer to DNA encapsidation while the latter are at an earlier stage in nucleocapsid synthesis. Taken together, these subnanometer resolution map observations strongly support the concept that CVSC occupancy can be used as an indicator for the progression of herpesvirus capsid assembly *in situ* and identifies capsids with varying levels of residual scaffold protein as CAIs along a continuum, having the potential for genome packaging rather than failed assembly dead ends.

### Portal vertex architecture differences between CAIs and C-capsids

In contrast to previous high-resolution single particle cryo-EM structures of biochemically purified herpesvirus capsids, the portal vertex for VZV capsids could not be reconstructed in previous reports ^39,41^ but has been achieved by Cao et al ^68^; this was not performed in infected cells *in situ*. To address this issue, we developed a RELION pipeline to reconstruct the VZV unique portal vertex, which was successfully identified for CAIs without the packaged dsDNA genome, or C-capsids within a subset of cryo-FIB/ET tomograms using STA and 3D classification of capsid vertices (**Fig. 6; Suppl. Fig. 16 to 18**). An initial cryo-ET map (23.6Å) for the unique portal vertex revealed a density within the capsid cavity reminiscent of the herpesvirus portal ^34,38,48–50^, and an absence of density for penton proteins compared to the 8.3Å cryo-ET map of the capsid vertex (**Fig. 3 to 5; Suppl. Fig. 17a**). Further STA of the portal density generated a 22.9Å cryo-ET map of the VZV portal with C12 symmetry (**Suppl. Fig. 17b**). A molecular model based on an AlphaFold prediction of an individual VZV portal protein (pORF54; 769aa) and modelling of 12 subunits based on the entire HSV-1 portal was placed into the 22.9Å cryo-ET map of the VZV portal (**Suppl. Fig. 17c and 17d**). Additional densities in the portal cryo-ET map were attributed to poor modelling by AlphaFold of amino acids 1 to 64 and 656 to 769, which were disordered and removed from the model. Architectural differences between the portal vertices of CAIs and C-capsids were visible when cryo-ET map reconstructions were performed based on capsid type (**Fig 6; Suppl. Fig. 18**). The portal cap was absent for CAIs compared to C-capsids, which was apparent when the entire capsid was reconstructed (**Suppl. Fig. 18**). The unique portal structure of C-capsids was confirmed as the portal density in the capsid cavity was only present at a single vertex (**Suppl. Fig. 18**). Further capsid-type specific architectural differences were evident from portal vertex cryo-ET maps and fitment of the molecular models for the peripentonal hexons, Ta and Tc triplexes, CVC1, ORF22 helices, and the portal (**Fig. 6**). As expected, densities for the pentonal MCP and alpha helices for the CVSC were absent and a channel for dsDNA translocation was visible through the center for the CAIs capsid portal vertex cryo-ET map (**Fig. 6b and 6c**). In contrast, the C-capsid cryo-ET map had a portal cap with densities for the CVSC, and which filled the portal vertex channel (**Fig 6e and 6f**), presumably, tentacle helices and the dsDNA genome terminus as previously described for HSV-1 ^38^. As expected from the VZV vertex focused classification analysis (**Fig. 5g to 5i**), occupancy of the CVSC at the non-portal vertices of the CAI was incomplete whereas each vertex for the C-capsid exhibited a full complement of the CVSC (**Fig. 6a and 6d; Suppl. Fig. 16**). These findings further support the CVSC as a defining element and thus a marker of herpesvirus capsid assembly and maturity inside the cell nucleus during replication.

**Figure 6.**
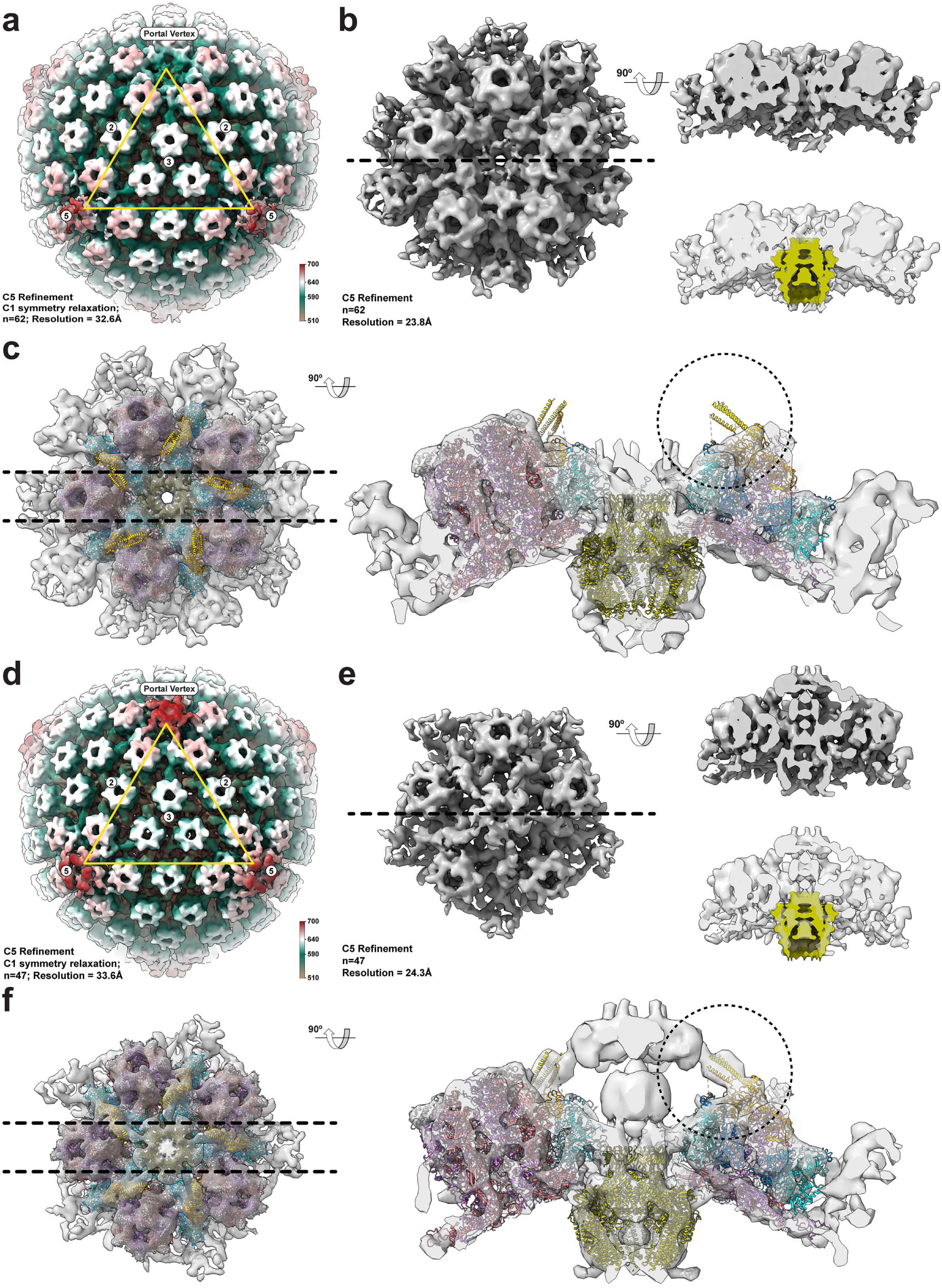
Differences in portal vertex architecture between VZV CAIs and C-capsids. **(a to f)** Cryo-ET maps of CAI and C-capsids and their portal vertices. **(a and d)** Cryo-ET maps of the VZV CAIs (A; EMD-49470) and C-capsid (D; EMD-49472) projected along the 3-fold axis of symmetry and colored radially (510-700Å form the center of the capsid). The yellow triangle encompasses the 2-, 3-, and 5-fold axes of symmetry. **(b and e)** Cryo-ET maps of the portal vertices from CAIs (B; EMD-49471) and C-capsids (E; EMD-49473). The left-hand panels show the vertices along the Z-axis of the cryo-ET map. The dashed black line shows the slice for views along the Y-axis, upper right-hand panels. The lower right-hand panels for the CAIs and C-capsids show transparent views of the cryo-ET maps with the portal map (see Suppl. Fig 15) aligned to the portal vertex map density. **(c and f)** The molecular model of the VZV capsid vertex without the pentonal MCP was fit into the cryo-ET maps of the CAIs and C-capsid portal vertices. The MCP of peripentonal hexons, triplexes Ta and Tc (TRX1, TRX2.1, and TRX2.2), small capsomere interacting protein (SCP), CVSC subunits CVC1, VP1/2.1 and VP1/2.2, and the VZV portal protein ORF54 are colored light coral or very soft violet (MCP), deep sky blue (TRX1), Columbia blue (TRX2.1), aqua (TRX2.2), dark gray (SCP), orange (CVC1), yellow (VP1/2.1), gold (VP1/2.2), and olive yellow (ORF54). Dashed lines indicate the slices used for the right-hand panels to more clearly visualize the portal. The dashed circles highlight the absence of CVSC densities in the portal vertex cryo-ET map for the B-capsid compared to the C-capsid (right-hand panels).

## DISCUSSION

The cell nucleus is a complex, crowded, dynamic environment, raising many questions about how herpesvirus capsids assemble within this large organelle. This study reveals the power of cryo-ET for the *in situ* investigation of herpesvirus assembly and morphogenesis at subnanometer resolution, the localization of macromolecular structures inside herpesvirus infected cell nuclei, and the reconstruction and segmentation of individual capsid proteins, allowing the identification of capsid assembly intermediates (CAIs) in their native environment for replication. In addition, the observations of procapsid precursors and architectural differences in capsid vertices revealed by the CVSC distribution on capsids in VZV infected cells provided new information relevant for understanding the temporal nature of herpesvirus capsid assembly inside the cell nucleus.

Herpesviruses are thought to assemble in a similar fashion to dsDNA bacteriophages where the capsomeres coalesce with the scaffold protein to generate procapsids ^28,29^. The high-resolution imaging inside the nucleus of VZV infected cells in the present study revealed angular segment structures where capsomeres, formed by the MCP and SCP, were associated with electron dense material presumed to be scaffold protein (**Suppl. Fig. 2**). This interpretation is supported by cell free systems that recapitulate herpesvirus capsid assembly by expression of the MCP, TRX1, TRX2 and VP22a proteins ^83^. VP22a is the scaffold protein expressed from UL26.5 (VZV ORF33.5) cleaved by the VP24 protease during maturation ^84,85^. VP24 is a cleavage product derived from UL26 (VZV ORF33), where the protease autocatalytically cleaves the UL26 gene product into VP24 and VP21. VP24 protease activity triggers spherical procapsids to undergo angularization and the SCP (VP26) is thought to provide additional capsid stability ^33,86–88^. For VZV, the ORF33 protease is required for replication and a SCP (ORF23) null virus produces predominantly spherical capsids, replicates poorly in cell culture and is non-viable in human skin xenografts ^89,90^. Furthermore, previous transient transfection studies have shown that the scaffold protein is required for intranuclear localization of the HSV-1 MCP (VP5) ^91^ whereas nuclear translocation of the VZV MCP is impaired by deleting the SCP ^90^. Further characterization of HSV procapsid formation in cell free expression systems noted the fragility and production of angular segments, leading to a proposed model where capsomeres and scaffold protein aggregate to form angular segments which in turn form spherical procapsids ^92^. In support of this, the angular segment structures of VZV demonstrate that herpesvirus procapsids form within the nucleus through cooperative assembly of capsomere-associated scaffold protein subunits rather than directly on a spherical precursor structure that recruits individual capsomeres.

Our study demonstrates that the acquisition of high-resolution 3D structure information derived from vitrified cells is critical for differentiating between capsid types that occur during assembly. Empty capsids were not apparent in the infected cell nucleus whereas previous observations with traditional thin section EM of infected cell nuclei frequently identify capsid cavities devoid of protein and dsDNA ^30^, leading to their conventional classification as A-capsids. From our 3D observations, evidence of scaffold remnants inside the capsid cavity were lost when the 2D sectional plane spans the entire diameter of the capsid, leading to their potential mischaracterization. Furthermore, our data eliminate fixation, sectioning, and staining artefacts, which could also lead to mischaracterization of A-capsids. In addition, capsid purification methods can produce empty capsids as an artefact since the pressurized dsDNA genome can be ejected due to the instability of the CVSC at the portal vertex ^32^. In contrast, the 3D analyses of capsid architecture coupled with the binding of CVSC heteropentamers to icosahedral vertices in the infected cell points to the need for an amended classification. In the present study, capsids with packaged dsDNA had the highest level of CVSC occupancy, consistent with previous single particle cryo-EM of purified C-capsids ^33,37,40,42,47^ and, furthermore, assembly intermediates were identifiable by correlating CVSC occupancy with levels of residual scaffold protein. Increasing CVSC occupancy at the vertices and of heteropentamer copies (up to the maximum of five) was associated with diminished quantities of scaffold protein in the capsid cavity. These observations indicate that conventionally designated B-capsids represent a continuum of CAIs that can potentially progress towards DNA encapsidation and formation of mature capsids ready for nuclear egress.

Earlier high-resolution cryo-EM studies of VZV capsids were not able to visualize the unique portal vertex ^39,41^. However, a recent study reconstructed the portal vertex from purified capsids ^68^. In the present *in situ* study, reconstruction of the portal vertex for CAIs and C-capsids along with acquisition of the VZV portal cap and packaged DNA for C-capsids was performed with capsids inside the infected cell nucleus. The dodecameric portal for VZV is composed of pORF54, which is essential for the packaging and cleavage of the dsDNA genome^93,94^. Similar to the VZV cryo-ET maps of the unique portal vertex in the present study, cryo-EM studies of purified HSV, HCMV, KSHV, and EBV have described the molecular architecture of the portal and portal vertex to have a C12 to C5 symmetry mismatch of the portal and vertex constituents ^34,35,38,49^. For HSV, the CVSC is visible at the portal vertex and the CVC2 protein is required for dsDNA packaging ^38^. The CVC1 subunit sits atop the Ta and Tc triplexes. At the portal, for HSV-1, the Ta triplexes are rotated 120° anticlockwise about their respective centers compared to pentonal Ta triplexes ^38^. Thus, the triplexes and the CVC1 are at an altered angle compared to the other 11 vertices. The MCP of HSV in the peripentonal hexon at position 6 has spine helices that extend into the pentonal space at non-portal vertices ^38^. The extension of the N-terminal helix was not visible in previous VZV cryo-EM structures, and that absence can now be attributed to the loss of ‘signal’ due to the combined STA of the portal and non-portal vertices ^39,41^. Moreover, the head groups of the CVC2 dimer, important for dsDNA genome packaging and retention, form part of the portal cap, which was not observed in previous studies of VZV, and explains the role of the CVSC in capsid maturation.

In addition to the retention of the dsDNA genome, the CVSC has also been linked to nuclear egress, a critical mechanism enabling capsids to leave the nucleus and enter the cytoplasm to complete herpesvirus morphogenesis ^95,96^. The process of nuclear egress is thought to be initiated by capsids budding at the nuclear envelope into the perinuclear space through interactions of CVC2 head groups with the nuclear egress complex (NEC) embedded in the INM, leading to acquisition of a primary envelope ^95,97^. Although the NEC has been shown to induce curvature in the absence of CVC2 via a mechanism that involves a random lattice formation ^98^, these studies were performed using a US3 mutant that causes capsids to become trapped within the perinuclear space ^99^. It has also been proposed that the CVSC functions as a selective switch to preferentially allow the dsDNA genome-filled C-capsids to leave the nucleus ^45,96^. Previous studies of capsid structures at subnanometer or near atomic resolution were performed with capsids purified from the nuclei of infected cells or purified herpesvirus particles, providing conflicting information about the quantities of the CVSC associated with capsid vertices ^33,36–41,47^. For VZV, precise information about the occupancy of the CVSC was unknown, likely due to the purification of these particles by ultracentrifugation and detergent-based removal of the envelope and tegument from C-capsids ^39,41^. Purification approaches have limitations in defining the stages of acquisition of the CVSC on capsids. However, the extensive interactions among all five subunits of the heteropentamer imply that the CVSC components co-assemble in the nucleus and act together as a functional unit in binding the capsid vertices ^33^. The present study provides definitive proof that the CVSC can assemble as a heteropentamer and bind to capsids inside the nuclei of VZV infected cells. Moreover, the application of STA and focused classification to the capsids further enabled the determination of the CVSC occupancy at individual sites at the vertices in relationship to different capsid types, supporting the notion that capsids with variable levels of scaffold protein are intermediates. Studies of HCMV support this concept, as the frequency of morphologically defined A-capsids within infected cell nuclei was very low compared to B- and C-capsids ^31,32^ and C-capsids purified from infected cell nuclei ejected their genomes leading to higher quantities of apparent A-capsids compared to those seen by TEM ^32^. Furthermore, the A-capsids were covered with the HCMV tegument protein pp150, which is equivalent to the CVSC of alphaherpesviruses and typically associated with C-capsids ^32,42,100^. Thus, high-resolution imaging coupled with state-of-the-art image classification approaches can now assign temporal information to the dynamic herpesvirus capsid assembly process *in situ*. The near complete occupancy of the CVSC on C-capsids and the increased levels of the CVSC on capsids with limited scaffold protein further support a selective function of the CVSC for nuclear egress, as both of these capsid types are found in extracellular VZV particles.

Our study provides direct evidence that the CVSC has a critical role in herpesvirus capsid assembly within the host cell nucleus, revealing heterogeneity in CVSC occupancy associated with CAIs, with the highest frequency found on dsDNA genome-filled C-capsids. Rather than discrete A-, B-, and C-capsid categories, we propose a model of herpesvirus capsid assembly where CVSC acquisition serves as a temporal marker towards completion; CAIs progress along a continuum defined by decreasing scaffold protein and increasing CVSC occupancy prior to genome packaging. Early intermediates retain abundant scaffold and have few CVSCs, whereas later intermediates contain limited scaffold with more extensive CVSC occupancy at the vertices, culminating in C-capsids with full vertex occupancy and packaged genomes. This model integrates structural heterogeneity into a dynamic pathway, resolves the ambiguity surrounding A-and B-capsids, and highlights CVSC acquisition as a critical determinant of capsid stability, genome retention, and readiness for nuclear egress. Our observations shed light on the unresolved question about whether A- and B-capsids are assembly dead ends, instead suggesting that A-capsids represent artefacts from purification whereas those conventionally grouped as B-capsids are assembly intermediates that can be further differentiated based on CVSC occupancy. Our findings underscore the intricate and dynamic nature of capsid assembly *in situ* during herpesvirus replication, providing new insights for the development of novel therapeutic strategies targeting herpesvirus capsid maturation.

## MATERIALS and METHODS

### Reagents, consumables, and resources

All reagents, consumables, and resources, including supplier information where applicable, are provided in **Suppl. Table 5**.

### Cells lines

MeWo cells (HTB-65; ATCC) were propagated in minimal essential medium (Corning Cellgro) supplemented with 10% fetal bovine serum (FBS; Invitrogen), nonessential amino acids (100 μM; Corning Cellgro), antibiotics (penicillin, 100 U/ml; streptomycin, 100 μg/ml; Invitrogen) and the antifungal agent amphotericin B (250 µg/ml; Invitrogen) at 37°C in a humidified atmosphere with 5% CO_2_.

### Viruses

The VZV parental Oka strain was originally cloned into a bacterial artificial chromosome (BAC) and designated pPOKA-BAC-DX ^101^. Recombinant viruses pOka-TK-GFP (pOka-rTK) and pOka-TK-RFP were generated in a previous study ^71,102^.

### ECM micropatterning of SiO_2_ EM grids

To position MeWo cells centrally within EM grid squares, 300 mesh gold grids coated with a thin perforated layer of SiO_2_ (Quantifoil R2/2) were subjected to micropatterning following the procedure available at protocols.io (https://www.protocols.io/view/micropatterning-em-grids-for-cryo-electron-tomogra-rm7vz3wqrgx1/v1). Briefly, EM grids were exposed to atmospheric plasma, incubated in poly-_L_-lysine (0.01%; Sigma) and subsequent incubations with 0.1M HEPES + mPEGSuccinimidyl Valerate (PEG-SVA MW 5,000; 100 mg/ml; Laysan Bio Inc.) then PLPP gel (1:6 dilution in H_2_O; Alveole). The coated grids were exposed to UV light according to a digitally generated micropattern (30×30µm or 33×33µm to match the 300 mesh spacing) that covered the EM grid square using a Primo photo-micropatterning system (Alveole) to selectively degrade the PEG coating to remove the hydrophobic antifouling layer. An extracellular matrix (ECM) surrogate, Gelatin Oregon Green 488 (1 mg/ml; Invitrogen), was applied to the grids prior to cell seeding.

### Vitrification of VZV Infected MeWo cells on ECM micropatterned grids

Cell free VZV cannot be purified to high titers making direct infection of cells grown on EM grids with high MOIs needed for productive infection impractical. Thus, infected cells were seeded directly onto ECM micropatterned EM grids. VZV (pOKA-TK-RFP) infected MeWo cells were passaged onto freshly seeded uninfected MeWo cells 24 hours prior to the day of seeding. On the day of seeding, the VZV infected MeWo cells were then passaged onto uninfected MeWo cells and incubated for 8 hours. The infected cells were then trypsinized, centrifuged at 400RCF for 5 minutes then resuspended in maintenance media to yield one cell per grid square; determined empirically on the day of seeding. Infected and uninfected cell suspensions were applied to the ECM micropatterned grids in 35mm glass bottomed dishes (MatTech P35G-1.5-14-C). After 2 hours incubation at 37°C in a humidified atmosphere with 5% CO_2,_ 3 ml of maintenance media was added and incubated for an additional 16 hours. Grids were removed from the maintenance media and plunge frozen into liquid ethane using an EM GP2 (Leica) with humidity at 99% and a blot time of 9 seconds.

### Cryo-FIB/SEM of vitrified VZV infected MeWo cells

An Aquilos 2 cryo-FIB/SEM microscope (ThermoFisher) was used to generate lamella from VZV infected cells on micropatterned grids. Prior to milling, the grids were platinum coated. Microexpansion joints of 1µm width and 4µm between the milled area and the microexpansion joint were used to relieve tension on the final lamellae^103^. The primary milling step was performed at 1nA and 5µm height and 15µm width with two 1µm microexpansion joints milled either side at 4 µm from the lamella milling site. Successive milling steps of 3µm (300pA), 1µm (100pA), 500nm (50pA), and a final polishing step of 175nm (30pA) were performed to generate lamella with a target thickness of 175nm. Lamella were produced for 6 to 10 cells per grid.

### Cryo-ET of whole or cryo-FIB/SEM milled VZV infected MeWo cells

Tilt series were either collected at the periphery of whole vitrified VZV infected MeWo cells on EM grids, using a 200kV Talos Arctica (FEI) or at cryo-FIB/SEM generated lamella, using a 300kV Titan Krios (FEI) controlled by SerialEM ^104^ to automate the data collection procedure using the parameters outlined in **Suppl. Table 1**. To identify areas of interest for tilt series collection, medium magnification (4,800-11,500x) montages were generated, and imaging sites were determined empirically by eye. Due to the large size of herpesvirus capsids, a 3.44Å/pixel size was chosen to accommodate areas where high frequencies of VZV capsids were present in the nucleus. A total of 399 tilt series were collected from cryo-FIB/SEM milled lamella across five, 3-day sessions on 300kV Titan Krios microscopes. A total of 194 tomograms were generated for VZV capsid quantification, analysis, and cryo-ET map reconstructions.

### Cryo-ET image processing

Motion correction, damage compensation, CTF estimation, and tilt series generation was performed using Warp ^105^. Tomogram reconstruction for visualization was performed using the IMOD tomography GUI Etomo ^106^. Tomogram processing and associated movie files were generated using 3dmod ^107^. Segmentation of tomograms was performed using Dragonfly software (Version 2022.2 for [Windows]. Comet Technologies Canada Inc., Montreal, Canada; software available at https://www.theobjects.com/dragonfly). Movies were generated using FFmpeg (FFmpeg Developers. (2016). ffmpeg tool (Version be1d324) available from http://ffmpeg.org/).

### Subnanometer resolution cryo-ET map reconstruction of the VZV capsid vertex

To reconstruct the VZV capsid vertex and 3-fold axis of symmetry a WARP-EMAN2 data processing strategy was used (**Suppl. Fig. 11**). Tilt series generated by WARP were imported into EMAN2 for tomogram alignment using patch tracking where handedness was checked and CTF measured. The convolutional neural network was employed to select particles (box size 512; n=1,084 capsids) from 194 tomograms and make sets (bin2). An initial model was generated from the sets using e2spt_sgd_new.py (--res=50.0; --niter=30; --shrink=4; --sym=icos) followed by subtomogram-subtilt refinement using e2spt_refine_new.py (--startres=30.0; --sym=icos; --iters=p3,t2,p,t,r,d; --keep=0.95; --tophat localwiener –goldstandard). Classification with icosahedral symmetry was performed on the output to remove poor quality particles that can negatively influence the subtomogram averaging; e2spt_refinemulti_new.py (--nref 3; --maxres 40; --loadali3d; --skipali; --sym icos; --maskalign spt_01/mask.hdf; --niter 20). Further refinement of 317 capsids produced a cryo-ET map of the entire VZV capsids of 13.0Å resolution. To generate a subnanometer resolution cryo-ET map for the VZV capsid vertex, symmetry expansion was performed with the 317 capsids using e2spt_extract.py (--postxf icos,0,0,110; --compressbits 8; --boxsz_unbin 192; --newlabel pent2; --mindist 500; --loadali2d) followed by building sets; e2spt_buildsets.py (--label=pent2; --allparticles; --spliteo). An average of the extracted particles was generated from the set using e2proc3d.py (--average; --averager mean.tomo; --avg_byxf). The average was then refined using e2spt_refine_new.py (--startres=30.0; --sym=c5; --iters=p3,t2,p,t,r,d; --keep=0.8,0.95,0.95; --tophat localwiener; --localrefine; --goldstandard) to achieve a cryo-ET map of the VZV capsid vertex at 8.3Å resolution. A similar procedure was used for the 3-fold axis of symmetry to achieve a cryo-ET map at 8.4Å resolution.

### Segmentation of the 3-fold and 5-fold cryo-ET maps

The 3- and 5-fold cryo-ET maps of the VZV capsid were segmented based on the 7BW6 asymmetric unit model of the VZV A-capsid ^39^ using ChimeraX ^108^. To segment the CVSC density in the 5-fold axis cryo-ET map, AlphaFold 2 ^109^ was used to model the heteropentamer based on the full-length proteins CVC1 (ORF43; monomer) and CVC2 (ORF34; dimer), and 45 C-terminal amino acids (2724-2768) of VP1/2 (ORF22). Molecular dynamics flexible fitting (MDFF) was performed to fit the two head groups of the CVC2 dimer into the cryo-ET map using NAMD ^110^ following the procedure outlined in the VMD tutorial. Conversion of the cryo-EM map to an MDFF potential, rigid body docking of the CVSC, initial structure generation, and secondary structure restraint definitions were all performed using VMD. The output files were used as input to run NAMD with the following parameters; step 1 -gscale 0.3, -numsteps 50000; step 2 -gscale 10, -numsteps 100000. A model of the asymmetric unit of the VZV capsid vertex, including the CVSC, was refined using ISOLDE ^111^. All images and movie files were generated using ChimeraX. Movies were generated using FFmpeg.

### Three axis orthogonal projection and uniform manifold approximation projection

The 3D VZV capsid particles were aligned to the symmetrized capsid reference from the EMAN2 icosahedral refinement. The subsequent 2D sub-tilt particles at the corresponding orientation were reconstructed with icosahedral symmetry to obtain symmetrized 3D particles free from missing wedge artifacts. Each particle was projected along three orthogonal axes of thin central slabs (72Å), using only one quarter of the volume closest to the center perpendicular to the corresponding axis. The resulting three 2D image projections per capsid were stacked and flattened into a 1D array that was used as input for uniform manifold approximation projection (UMAP). After manifold embedding, particles on the 2D UMAP space were clustered using a K-means algorithm to produce the individual classes.

### Reconstruction of the VZV Portal Vertex

To reconstruct the VZV portal vertex a WARP-IMOD-EMAN2-Relion data processing strategy was used (**Suppl. Fig. 16**). Tilt series were aligned using IMOD and for the final aligned stack, unbinned, CTF estimation was performed using CTFplotter with the ‘Find astigmatism option’ checked. Tomograms for Relion were generated using relion_tomo_reconstruct_tomogram and used for particle picking of capsids in EMAN2 (n=471). Tomograms and particle coordinates were imported into Relion using Tomo Import. Make pseudo-subtomos was used to make subtomograms (bin 4; 13.76 Å/pixel) using a box size (pixels) of 360 and cropped box size (pixels) of 180. An initial model was generated using 3D initial model and a mask was generated using Dynamo and Mask create. A 3D refinement was performed with 3D auto-refine with an initial low-pass filter of 50 Å and I3 symmetry. Particle reconstruction was performed (bin2; 6.88Å/pixel) with Tomo reconstruct particle with a box size of 440 and cropped box size of 220 with I3 symmetry, which was used to generate a new mask, followed by reconstruction of subtomograms using Make pseudo-subtomos. A 3D refinement was performed with 3D auto-refine with an initial low-pass filter of 30Å and I3 symmetry. The output was used for Tomo frame alignment, particle reconstruction (bin 1; 3.44Å/pixel; box size 440 pixels; I3 symmetry), mask creation, reconstruction of subtomograms, and 3D refinement with an initial low-pass filter of 14Å and I3 symmetry, generating a 12.1Å cryo-ET map of the VZV capsid.

Symmetry expansion was performed using relion_particle_symmetry_expand with --sym I3. This process produces 5 rotations per vertex; redundant _rlnAngleRot values were removed to leave 12 unique vertices per capsid (n=5,652). Vertices beyond the boundaries of the tomograms were removed (n=5,318). The Z height was adjusted to center the vertices in the center of the box using relion_star_handler and --center_Z command. The output was used for particle reconstruction (bin 4; 13.76 Å/pixel; initial box size 120 pixels; cropped box size 60 pixels; C5 symmetry), mask creation, reconstruction of subtomograms, and 3D refinement with an initial low-pass filter of 50Å and C5 symmetry. The output was used for particle reconstruction (bin 2; 6.88 Å/pixel; box size 240 pixels; cropped box size 120 pixels; C5 symmetry), mask creation, reconstruction of subtomograms, and 3D refinement with an initial low-pass filter of 30Å and C5 symmetry. The output was used for particle reconstruction (bin 1; 3.44Å/pixel; box size 240 pixels; C5 symmetry), mask creation, reconstruction of subtomograms, and 3D refinement with an initial low-pass filter of 15Å and C5 symmetry, generating a 10.3Å cryo-ET map of the VZV capsid vertex.

The output was used to generate a cryo-ET capsid vertex map at bin 4 using the above processing scheme then used to perform 3D classification (12 classes; 29 iterations). An initial 60 particles were selected based on the presence of a portal density and the absence of the pentonal major capsid protein then used to generate a cryo-ET portal vertex map at bin 4 using the above processing scheme. A focused cylindrical mask was generated and used for further 3D classification of the VZV capsid vertices yielding a total of 103 portal vertices. A final round of template-based 3D classification yielded a total of 109 portal vertices. The output was used for particle reconstruction (bin 2; 6.88Å/pixel; box size 240 pixels; cropped box size 120 pixels; C5 symmetry), mask creation, reconstruction of subtomograms, and 3D refinement with an initial low-pass filter of 30Å and C5 symmetry, generating a 23.6Å cryo-ET map of the VZV capsid portal vertex. To reconstruct the portal, particle reconstruction (bin 2; 6.88Å/pixel; box size 90 pixels; cropped box size 30 pixels; C12 symmetry), mask creation, reconstruction of subtomograms, and 3D refinement with an initial low-pass filter of 30Å, C12 symmetry, and C1 symmetry relaxation was performed, generating a 22.9Å cryo-ET map of the VZV portal.

To identify portal vertex specific features, the 109 portal vertices were divided into CAI (n=62) and C-capsids (n=47) based on morphological features; either scaffold protein (CAI) or genomic DNA (C-capsids) inside the capsid cavity. Cryo-ET maps for the portal vertices from CAI (23.8Å) and C-capsids (24.3Å) were generated as described above; particle reconstruction (bin 2; 6.88Å/pixel; box size 240 pixels; cropped box size 120 pixels; C5 symmetry), mask creation, reconstruction of subtomograms, and 3D refinement with an initial low-pass filter of 30Å and C5 symmetry. Cryo-ET maps for the entire capsid from CAI (32.6Å) and C-capsids (33.6Å) were generated by first adjusting the Z-height of the particles as previously described the performing particle reconstruction (bin 2; 6.88Å/pixel; box size 512 pixels; cropped box size 256 pixels; I3 symmetry), mask creation, reconstruction of subtomograms, and 3D refinement with an initial low-pass filter of 30Å, C5 symmetry, and C1 symmetry relaxation.

The resolution of all Relion generated cryo-ET maps were confirmed using post-processing with soft masks and assessed by Fourier Shell correlation curves with a cut-off at 0.143 from two independent half-sets of the pseudosubtomograms.

### Statistics and Reproducibility

All quantitative data were analyzed with two-way ANOVA to determine statistical significance using Prism (GraphPad Software).

## Supporting information

Movie Titles

## DATA AVAILABILITY

Data generated and/or analyzed during the current study are available in the paper, appended as supplementary data, available in public repositories. The original cryo-ET movie files, tilt series, and tomograms have been deposited in the Electron Microscopy Public Image Archive (EMPIAR) with accession code EMPIAR-12464. The cryo-ET maps have been deposited in the Electron Microscopy Data Bank (EMDB) with accession codes EMD-49591 for the vertex cryo-ET map (model PDB 9NO1), EMD-49886 for the 3-fold axis cryo-ET map and EMD-49465 to 49473 for cryo-ET maps of the portal vertex. The source data underlying Figures 5A, 5E, and 5I are provided in the associated Source Data file.

## ACKNOWLEDGMENTS

Cryo-FIB-SEM and cryo-ET were performed at the Stanford-SLAC Cryo-EM Center and the Stanford-SLAC Cryo-ET Specimen Preparation Center under the support of the National Institutes of Health Common Fund’s Transformative High Resolution Cryo-electron Microscopy Program (U24GM139166 and U24GM129541); and the NIH Shared Instrumentation Grant (S10OD021600). Research support was provided by NIH grants R01AI020459 (SLO), R21AI159375 (SLO and WC), R01GM150905 (MC), AI15937502 (WC), and R21MH125285 (MC). Molecular graphics and analyses performed with UCSF ChimeraX, developed by the Resource for Biocomputing, Visualization, and Informatics at the University of California, San Francisco, with support from National Institutes of Health R01GM129325 and the Office of Cyber Infrastructure and Computational Biology, National Institute of Allergy and Infectious Diseases.

## AUTHOR CONTRIBUTIONS

SLO: Conceptualization, methodology, software application, validation, formal analysis, investigation, resources, data curation, writing - original draft and editing, visualization, supervision, project administration, funding acquisition. MC: Software application, validation, formal analysis, writing - review and editing. LE: Methodology, resources, writing - review and editing. CWH: Methodology, writing - review and editing. XZ: Methodology, writing - review and editing. MFS: Validation, formal analysis, writing - review and editing. AMA: Resources, writing - review and editing, supervision, funding acquisition. WC: Resources, writing - review and editing, supervision, funding acquisition.

## REFERENCES

1 Arvin, A. M. & Abendroth, A. in Fields Virology: DNA Viruses, 7e (eds Peter M. Howley, David M. Knipe, Blossom A. Damania, & Jeffrey I. Cohen) 0 (Lippincott Williams & Wilkins, a Wolters Kluwer business, 2022).

2 Damania, B. & Cesarman, E. in Fields Virology: DNA Viruses, 7e (eds Peter M. Howley, David M. Knipe, Blossom A. Damania, & Jeffrey I. Cohen) 0 (Lippincott Williams & Wilkins, a Wolters Kluwer business, 2022).

3 Gewurz, B. E., Longnecker, R. M. & Cohen, J. I. in Fields Virology: DNA Viruses, 7e (eds Peter M. Howley, David M. Knipe, Blossom A. Damania, & Jeffrey I. Cohen) 0 (Lippincott Williams & Wilkins, a Wolters Kluwer business, 2022).

4 Goodrum, F., Britt, W. & Mocarski, E. S. in Fields Virology: DNA Viruses, 7e (eds Peter M. Howley, David M. Knipe, Blossom A. Damania, & Jeffrey I. Cohen) 0 (Lippincott Williams & Wilkins, a Wolters Kluwer business, 2022).

5 Knipe, D. M., Heldwein, E. E., Mohr, I. J. & Sodroski, C. N. in Fields Virology: DNA Viruses, 7e (eds Peter M. Howley, David M. Knipe, Blossom A. Damania, & Jeffrey I. Cohen) 0 (Lippincott Williams & Wilkins, a Wolters Kluwer business, 2022).

6 Krug, L. T. & Pellett, P. E. in Fields Virology: DNA Viruses, 7e (eds Peter M. Howley, David M. Knipe, Blossom A. Damania, & Jeffrey I. Cohen) 0 (Lippincott Williams & Wilkins, a Wolters Kluwer business, 2022).

7 Mori, Y., Zerr, D. M., Flamand, L. & Pellett, P. E. in Fields Virology: DNA Viruses, 7e (eds Peter M. Howley, David M. Knipe, Blossom A. Damania, & Jeffrey I. Cohen) 0 (Lippincott Williams & Wilkins, a Wolters Kluwer business, 2022).

8 Whitley, R. J. & Johnston, C. in Fields Virology: DNA Viruses, 7e (eds Peter M. Howley, David M. Knipe, Blossom A. Damania, & Jeffrey I. Cohen) 0 (Lippincott Williams & Wilkins, a Wolters Kluwer business, 2022).

9 Gilden, D. H., Mahalingham, R., Deitch, S. & Randall, C., J. in Alpha herpesviruses : molecular and cellular biology (ed R. M. Sandri-Goldin) 402 (Caister Academic, 2006).

10 Banovic, T. et al. Disseminated varicella infection caused by varicella vaccine strain in a child with low invariant natural killer T cells and diminished CD1d expression. J Infect Dis 204, 1893–1901, (2011).

11 Gilden, D., Cohrs, R. J., Mahalingam, R. & Nagel, M. A. Varicella zoster virus vasculopathies: diverse clinical manifestations, laboratory features, pathogenesis, and treatment. Lancet Neurol 8, 731–740, (2009).

12 Jean-Philippe, P. et al. Severe varicella caused by varicella-vaccine strain in a child with significant T-cell dysfunction. Pediatrics 120, e1345–1349, (2007).

13 Johnson, R. W. Zoster-associated pain: what is known, who is at risk and how can it be managed? Herpes 14 **Suppl 2**, 30–34, (2007).

14 Pahud, B. A., Glaser, C. A., Dekker, C. L., Arvin, A. M. & Schmid, D. S. Varicella zoster disease of the central nervous system: epidemiological, clinical, and laboratory features 10 years after the introduction of the varicella vaccine. J Infect Dis 203, 316–323, (2011).

15 Schmader, K. Herpes zoster and postherpetic neuralgia in older adults. Clin Geriatr Med 23, 615–632, vii-viii, (2007).

16 Brunell, P. A., Geiser, C. F., Novelli, V., Lipton, S. & Narkewicz, S. Varicella-like illness caused by live varicella vaccine in children with acute lymphocytic leukemia. Pediatrics 79, 922–927, (1987).

17 Marin, M., Meissner, H. C. & Seward, J. F. Varicella prevention in the United States: a review of successes and challenges. Pediatrics 122, e744–751, (2008).

18 Pickering, L. K. et al. Immunization programs for infants, children, adolescents, and adults: clinical practice guidelines by the Infectious Diseases Society of America. Clin Infect Dis 49, 817–840, (2009).

19 Han, J. Y., Hanson, D. C. & Way, S. S. Herpes zoster and meningitis due to reactivation of varicella vaccine virus in an immunocompetent child. Pediatr Infect Dis J 30, 266–268, (2011).

20 Krause, P. R. & Klinman, D. M. Varicella vaccination: evidence for frequent reactivation of the vaccine strain in healthy children. Nat Med 6, 451–454, (2000).

21 Whitley, R. J. Therapy of herpes virus infections in children. Adv Exp Med Biol 609, 216–232, (2008).

22 Whitley, R. J., Volpi, A., McKendrick, M., Wijck, A. & Oaklander, A. L. Management of herpes zoster and post-herpetic neuralgia now and in the future. J Clin Virol 48 **Suppl 1**, S20–28, (2010).

23 Shiraki, K., Takemoto, M. & Daikoku, T. Emergence of varicella-zoster virus resistance to acyclovir: epidemiology, prevention, and treatment. Expert Rev Anti Infect Ther 19, 1415–1425, (2021).

24 Topalis, D., Gillemot, S., Snoeck, R. & Andrei, G. Thymidine kinase and protein kinase in drug-resistant herpesviruses: Heads of a Lernaean Hydra. Drug Resist Updat 37, 1–16, (2018).

25 Harfouche, M. et al. Estimated global and regional incidence and prevalence of herpes simplex virus infections and genital ulcer disease in 2020: mathematical modelling analyses. Sex Transm Infect, (2024).

26 Chaiyakunapruk, N. et al. Estimated global and regional economic burden of genital herpes simplex virus infection among 15-49 year-olds in 2016. BMC Glob Public Health 2, 42, (2024).

27 Davison, A. J. in Human Herpesviruses: Biology, Therapy, and Immunoprophylaxis (eds A. Arvin et al.) (2007).

28 Baines, J. D. Herpes simplex virus capsid assembly and DNA packaging: a present and future antiviral drug target. Trends Microbiol 19, 606–613, (2011).

29 Heming, J. D., Conway, J. F. & Homa, F. L. Herpesvirus Capsid Assembly and DNA Packaging. Adv Anat Embryol Cell Biol 223, 119–142, (2017).

30 Gibson, W. & Roizman, B. Proteins specified by herpes simplex virus. 8. Characterization and composition of multiple capsid forms of subtypes 1 and 2. J Virol 10, 1044–1052, (1972).

31 Tandon, R., Mocarski, E. S. & Conway, J. F. The A, B, Cs of herpesvirus capsids. Viruses 7, 899–914, (2015).

32 Stevens, A., Cruz-Cosme, R., Armstrong, N., Tang, Q. & Zhou, Z. H. Structure-guided mutagenesis targeting interactions between pp150 tegument protein and small capsid protein identify five lethal and two live-attenuated HCMV mutants. Virology 596, 110115, (2024).

33 Dai, X. & Zhou, Z. H. Structure of the herpes simplex virus 1 capsid with associated tegument protein complexes. Science 360, (2018).

34 Gong, D. et al. DNA-Packing Portal and Capsid-Associated Tegument Complexes in the Tumor Herpesvirus KSHV. Cell 178, 1329–1343 e1312, (2019).

35 Li, Z., Pang, J., Dong, L. & Yu, X. Structural basis for genome packaging, retention, and ejection in human cytomegalovirus. Nat Commun 12, 4538, (2021).

36 Li, Z. et al. CryoEM structure of the tegumented capsid of Epstein-Barr virus. Cell Res 30, 873–884, (2020).

37 Liu, W. et al. Structures of capsid and capsid-associated tegument complex inside the Epstein-Barr virus. Nat Microbiol 5, 1285–1298, (2020).

38 Liu, Y. T., Jih, J., Dai, X., Bi, G. Q. & Zhou, Z. H. Cryo-EM structures of herpes simplex virus type 1 portal vertex and packaged genome. Nature 570, 257–261, (2019).

39 Sun, J. et al. Cryo-EM structure of the varicella-zoster virus A-capsid. Nat Commun 11, 4795, (2020).

40 Wang, G. et al. Structures of pseudorabies virus capsids. Nat Commun 13, 1533, (2022).

41 Wang, W., et al. Near-atomic cryo-electron microscopy structures of varicella-zoster virus capsids. Nat Microbiol 5, 1542–1552, (2020).

42 Yu, X., Jih, J., Jiang, J. & Zhou, Z. H. Atomic structure of the human cytomegalovirus capsid with its securing tegument layer of pp150. Science 356, (2017).

43 Zhang, Y. et al. Atomic structure of the human herpesvirus 6B capsid and capsid-associated tegument complexes. Nat Commun 10, 5346, (2019).

44 Dai, X. et al. Structure and mutagenesis reveal essential capsid protein interactions for KSHV replication. Nature 553, 521–525, (2018).

45 Huet, A., Huffman, J. B., Conway, J. F. & Homa, F. L. Role of the Herpes Simplex Virus CVSC Proteins at the Capsid Portal Vertex. J Virol 94, (2020).

46 Sae-Ueng, U. et al. Major capsid reinforcement by a minor protein in herpesviruses and phage. Nucleic Acids Res 42, 9096–9107, (2014).

47 Wang, J. et al. Structure of the herpes simplex virus type 2 C-capsid with capsid-vertex-specific component. Nat Commun 9, 3668, (2018).

48 Li, Z. et al. Cryo-electron microscopy structures of capsids and in situ portals of DNA-devoid capsids of human cytomegalovirus. Nat Commun 14, 2025, (2023).

49 Machon, C. et al. Atomic structure of the Epstein-Barr virus portal. Nat Commun 10, 3891, (2019).

50 Schmid, M. F. et al. A tail-like assembly at the portal vertex in intact herpes simplex type-1 virions. PLoS Pathog 8, e1002961, (2012).

51 Draganova, E. B., Valentin, J. & Heldwein, E. E. The Ins and Outs of Herpesviral Capsids: Divergent Structures and Assembly Mechanisms across the Three Subfamilies. Viruses 13, (2021).

52 Diwaker, D. & Wilson, D. W. Microtubule-Dependent Trafficking of Alphaherpesviruses in the Nervous System: The Ins and Outs. Viruses 11, (2019).

53 Zaichick, S. V. et al. The herpesvirus VP1/2 protein is an effector of dynein-mediated capsid transport and neuroinvasion. Cell Host Microbe 13, 193–203, (2013).

54 Jih, J., Liu, Y. T., Liu, W. & Zhou, Z. H. The incredible bulk: Human cytomegalovirus tegument architectures uncovered by AI-empowered cryo-EM. Sci Adv 10, eadj1640, (2024).

55 Buch, M. H. C., Newcomb, W. W., Winkler, D. C., Steven, A. C. & Heymann, J. B. Cryo-Electron Tomography of the Herpesvirus Procapsid Reveals Interactions of the Portal with the Scaffold and a Shift on Maturation. mBio 12, (2021).

56 Cardone, G. et al. Visualization of the herpes simplex virus portal in situ by cryo-electron tomography. Virology 361, 426–434, (2007).

57 Dai, W. et al. Unique structures in a tumor herpesvirus revealed by cryo-electron tomography and microscopy. J Struct Biol 161, 428–438, (2008).

58 Deng, B., O’Connor, C. M., Kedes, D. H. & Zhou, Z. H. Direct visualization of the putative portal in the Kaposi’s sarcoma-associated herpesvirus capsid by cryoelectron tomography. J Virol 81, 3640–3644, (2007).

59 Deng, B., O’Connor, C. M., Kedes, D. H. & Zhou, Z. H. Cryo-electron tomography of Kaposi’s sarcoma-associated herpesvirus capsids reveals dynamic scaffolding structures essential to capsid assembly and maturation. J Struct Biol 161, 419–427, (2008).

60 Grunewald, K. et al. Three-dimensional structure of herpes simplex virus from cryo-electron tomography. Science 302, 1396–1398, (2003).

61 Newcomb, W. W. et al. The Primary Enveloped Virion of Herpes Simplex Virus 1: Its Role in Nuclear Egress. mBio 8, (2017).

62 Si, Z. et al. Different functional states of fusion protein gB revealed on human cytomegalovirus by cryo electron tomography with Volta phase plate. PLoS Pathog 14, e1007452, (2018).

63 Chang, J. T., Schmid, M. F., Rixon, F. J. & Chiu, W. Electron cryotomography reveals the portal in the herpesvirus capsid. J Virol 81, 2065–2068, (2007).

64 Vijayakrishnan, S., McElwee, M., Loney, C., Rixon, F. & Bhella, D. In situ structure of virus capsids within cell nuclei by correlative light and cryo-electron tomography. Sci Rep 10, 17596, (2020).

65 Bucks, M. A., O’Regan, K. J., Murphy, M. A., Wills, J. W. & Courtney, R. J. Herpes simplex virus type 1 tegument proteins VP1/2 and UL37 are associated with intranuclear capsids. Virology 361, 316–324, (2007).

66 Fan, W. H. et al. The large tegument protein pUL36 is essential for formation of the capsid vertex-specific component at the capsid-tegument interface of herpes simplex virus 1. J Virol 89, 1502–1511, (2015).

67 Schipke, J. et al. The C terminus of the large tegument protein pUL36 contains multiple capsid binding sites that function differently during assembly and cell entry of herpes simplex virus. J Virol 86, 3682–3700, (2012).

68 Cao, L. et al. Insights into varicella-zoster virus assembly from the B- and C-capsid at near-atomic resolution structures. hLife 2, 64–74, (2024).

69 Reimer, L. & Kohl, H. Transmission electron microscopy : physics of image formation. Fifth edition. edn, (Springer, 2008).

70 Schaffer, M. et al. A cryo-FIB lift-out technique enables molecular-resolution cryo-ET within native Caenorhabditis elegans tissue. Nat Methods 16, 757–762, (2019).

71 Oliver, S. L., Yang, E. & Arvin, A. M. Dysregulated Glycoprotein B-Mediated Cell-Cell Fusion Disrupts Varicella-Zoster Virus and Host Gene Transcription during Infection. J Virol 91, (2017).

72 Oliver, S. L. et al. A glycoprotein B-neutralizing antibody structure at 2.8 A uncovers a critical domain for herpesvirus fusion initiation. Nat Commun 11, 4141, (2020).

73 Oliver, S. L. et al. The N-terminus of varicella-zoster virus glycoprotein B has a functional role in fusion. PLoS Pathog 17, e1008961, (2021).

74 Zhou, M., Kamarshi, V., Arvin, A. M. & Oliver, S. L. Calcineurin phosphatase activity regulates Varicella-Zoster Virus induced cell-cell fusion. PLoS Pathog 16, e1009022, (2020).

75 Baker, M. L., Jiang, W., Rixon, F. J. & Chiu, W. Common ancestry of herpesviruses and tailed DNA bacteriophages. J Virol 79, 14967–14970, (2005).

76 Homa, F. L. et al. Structure of the pseudorabies virus capsid: comparison with herpes simplex virus type 1 and differential binding of essential minor proteins. J Mol Biol 425, 3415–3428, (2013).

77 DeRussy, B. M., Boland, M. T. & Tandon, R. Human Cytomegalovirus pUL93 Links Nucleocapsid Maturation and Nuclear Egress. J Virol 90, 7109–7117, (2016).

78 Evilevitch, A. & Sae-Ueng, U. Mechanical Capsid Maturation Facilitates the Resolution of Conflicting Requirements for Herpesvirus Assembly. J Virol 96, e0183121, (2022).

79 McInnes, L. & Healy, J. UMAP: Uniform Manifold Approximation and Projection for Dimension Reduction. ArXiv abs/1802.03426, (2018).

80 Chen, M. & Ludtke, S. J. Deep learning-based mixed-dimensional Gaussian mixture model for characterizing variability in cryo-EM. Nat Methods 18, 930–936, (2021).

81 Zhong, E. D., Bepler, T., Berger, B. & Davis, J. H. CryoDRGN: reconstruction of heterogeneous cryo-EM structures using neural networks. Nat Methods 18, 176–185, (2021).

82 Radermacher, M. Three-dimensional reconstruction of single particles from random and nonrandom tilt series. J Electron Microsc Tech 9, 359–394, (1988).

83 Newcomb, W. W., Homa, F. L., Thomsen, D. R., Ye, Z. & Brown, J. C. Cell-free assembly of the herpes simplex virus capsid. J Virol 68, 6059–6063, (1994).

84 Davison, M. D., Rixon, F. J. & Davison, A. J. Identification of genes encoding two capsid proteins (VP24 and VP26) of herpes simplex virus type 1. J Gen Virol 73 **( Pt** **10**), 2709–2713, (1992).

85 Preston, V. G. et al. Efficient herpes simplex virus type 1 (HSV-1) capsid formation directed by the varicella-zoster virus scaffolding protein requires the carboxy-terminal sequences from the HSV-1 homologue. J Gen Virol 78 **( Pt** **7**), 1633–1646, (1997).

86 Yuan, S. et al. Cryo-EM structure of a herpesvirus capsid at 3.1 A. Science 360, (2018).

87 Zhou, Z. H. et al. Assembly of VP26 in herpes simplex virus-1 inferred from structures of wild-type and recombinant capsids. Nat Struct Biol 2, 1026–1030, (1995).

88 Cardone, G., Heymann, J. B., Cheng, N., Trus, B. L. & Steven, A. C. Procapsid assembly, maturation, nuclear exit: dynamic steps in the production of infectious herpesvirions. Adv Exp Med Biol 726, 423–439, (2012).

89 Zhang, Z. et al. Genome-wide mutagenesis reveals that ORF7 is a novel VZV skin-tropic factor. PLoS Pathog 6, e1000971, (2010).

90 Chaudhuri, V., Sommer, M., Rajamani, J., Zerboni, L. & Arvin, A. M. Functions of Varicella-zoster virus ORF23 capsid protein in viral replication and the pathogenesis of skin infection. J Virol 82, 10231–10246, (2008).

91 Nicholson, P. et al. Localization of the herpes simplex virus type 1 major capsid protein VP5 to the cell nucleus requires the abundant scaffolding protein VP22a. J Gen Virol 75 **( Pt** **5**), 1091–1099, (1994).

92 Newcomb, W. W. et al. Assembly of the herpes simplex virus capsid: characterization of intermediates observed during cell-free capsid formation. J Mol Biol 263, 432–446, (1996).

93 Visalli, M. A., House, B. L., Selariu, A., Zhu, H. & Visalli, R. J. The varicella-zoster virus portal protein is essential for cleavage and packaging of viral DNA. J Virol 88, 7973–7986, (2014).

94 Visalli, R. J. & Howard, A. J. Non-axial view of the varicella-zoster virus portal protein reveals conserved crown, wing and clip architecture. Intervirology 57, 121–125, (2014).

95 Draganova, E. B., Zhang, J., Zhou, Z. H. & Heldwein, E. E. Structural basis for capsid recruitment and coat formation during HSV-1 nuclear egress. Elife 9, (2020).

96 Yang, K. & Baines, J. D. Selection of HSV capsids for envelopment involves interaction between capsid surface components pUL31, pUL17, and pUL25. Proc Natl Acad Sci U S A 108, 14276–14281, (2011).

97 Mettenleiter, T. C., Muller, F., Granzow, H. & Klupp, B. G. The way out: what we know and do not know about herpesvirus nuclear egress. Cell Microbiol 15, 170–178, (2013).

98. Prazak, V., et al. Molecular plasticity of herpesvirus nuclear egress analysed in situ. Nat Microbiol 9, 1842-1855, (2024).

99 Klupp, B. G., Granzow, H. & Mettenleiter, T. C. Nuclear envelope breakdown can substitute for primary envelopment-mediated nuclear egress of herpesviruses. J Virol 85, 8285–8292, (2011).

100 Dai, X. et al. The smallest capsid protein mediates binding of the essential tegument protein pp150 to stabilize DNA-containing capsids in human cytomegalovirus. PLoS Pathog 9, e1003525, (2013).

101 Tischer, B. K. et al. A self-excisable infectious bacterial artificial chromosome clone of varicella-zoster virus allows analysis of the essential tegument protein encoded by ORF9. J Virol 81, 13200–13208, (2007).

102 Yang, E., Arvin, A. M. & Oliver, S. L. The cytoplasmic domain of varicella-zoster virus glycoprotein H regulates syncytia formation and skin pathogenesis. PLoS Pathog 10, e1004173, (2014).

103 Wolff, G. et al. Mind the gap: Micro-expansion joints drastically decrease the bending of FIB-milled cryo-lamellae. J Struct Biol 208, 107389, (2019).

104 Mastronarde, D. N. Automated electron microscope tomography using robust prediction of specimen movements. J Struct Biol 152, 36–51, (2005).

105 Tegunov, D. & Cramer, P. Real-time cryo-electron microscopy data preprocessing with Warp. Nat Methods 16, 1146–1152, (2019).

106 Mastronarde, D. N. Dual-axis tomography: an approach with alignment methods that preserve resolution. J Struct Biol 120, 343–352, (1997).

107 Kremer, J. R., Mastronarde, D. N. & McIntosh, J. R. Computer visualization of three-dimensional image data using IMOD. J Struct Biol 116, 71–76, (1996).

108 Pettersen, E. F. et al. UCSF ChimeraX: Structure visualization for researchers, educators, and developers. Protein Sci 30, 70–82, (2021).

109 Jumper, J. et al. Highly accurate protein structure prediction with AlphaFold. Nature 596, 583–589, (2021).

110 Phillips, J. C. et al. Scalable molecular dynamics on CPU and GPU architectures with NAMD. J Chem Phys 153, 044130, (2020).

111 Croll, T. I. ISOLDE: a physically realistic environment for model building into low-resolution electron-density maps. Acta Crystallogr D Struct Biol 74, 519–530, (2018).

