## Supplementary material for "Cryogenic Electron Tomography Redefines Herpesvirus Capsid Assembly Intermediates Inside the Cell Nucleus": Movie Titles

**Supplemental Movie 1.** **A Tomogram of a VZV Capsid Docked at a Nuclear Pore.** Of the 720 slices for the IMOD reconstructed tomogram (EMPIAR-12464; 20220507_CS02_VZV52G2_lamella3_tilt02.mrc_rec.mrc), the movie focusses on a VZV capsid at a nuclear pore and moves through slices 130 to 290 then back to slice 130. A single slice (179) is annotated in Fig. 1Ei and Suppl. Fig. 1. The white scale bar depicts 125nm.

**Supplemental Movie 2.** **A Tomogram of Intranuclear VZV Capsids.** Of the 528 slices for the IMOD reconstructed tomogram (EMPIAR-12464; 20211210_CS02_VZV46G2_Tilt19.mrc_rec.mrc), the movie focusses on a cluster of VZV capsids inside the nucleus of an infected cell and moves through slices 1 to 430 then back to slice 1. A single slice (243) is annotated in Fig. 1Eii and Suppl. Fig. 3. The white scale bar depicts 125nm.

**Supplemental Movie 3.** **A Tomogram of a VZV Capsid in the Process of Nuclear Egress.** Of the 646 slices for the IMOD reconstructed tomogram (EMPIAR-12464; 20220615-CS02_VZV51G2_lamella2_tilt11.mrc_rec.mrc), the movie focusses on a C-capsid in the process of nuclear egress from the infected cell nucleus and moves through slices 100 to 410 then back to slice 100. A single slice (253) is annotated in Fig. 1Eiii and Suppl. Fig. 4. The white scale bar depicts 125nm.

**Supplemental Movie 4.** **A Tomogram of a VZV Capsid in the Process of Secondary Envelopment.** Of the 676 slices for the IMOD reconstructed tomogram (EMPIAR-12464; 20220509_CS02_VZV51G1_lamella2_tilt03.mrc_rec.mrc), the movie focusses on a C-capsid in the process of secondary envelopment in the infected cell cytoplasm and moves through slices 100 to 340 then back to slice 100. A single slice (155) is annotated in Fig. 1Eiv and Suppl. Fig. 5. The white scale bar depicts 125nm.

**Supplemental Movie 5.** **A Tomogram of VZV Capsids in the Undergoing Morphogenesis.** Of the 450 slices for the IMOD reconstructed tomogram (EMPIAR-12464; 20220507_CS02_VZV52G2_lamella3_tilt03.mrc_rec.mrc), the movie focusses on VZV particle morphogenesis in the infected cell cytoplasm and moves through slices 60 to 300 then back to slice 60. A single slice (143) is annotated in Fig. 1Ev and Suppl. Fig. 6. The white scale bar depicts 125nm.

**Supplemental Movie 6.** **A Tomogram of Extracellular VZV Particles Bound to the Plasma Membrane.** Of the 475 slices for the IMOD reconstructed tomogram (EMPIAR-12464; 191017_VZV6_G4_11117b_Tilt28.mrc_rec.mrc), the movie focusses on extracellular VZV particles bound to the plasma membrane and moves through slices 1 to 220 then back to slice 1. Single slices (64 and 163) are annotated in Fig. 1Evi (64) and Suppl. Fig. 6 (64 and 163). The white scale bar depicts 125nm.

**Supplemental Movie 7.** **Segmentation of the VZV Capsid Vertex Cryo-ET Map.** The movie shows segmentation of the 8.3Å VZV capsid vertex cryo-ET map based on the 7BW6 model refined using the subnanometer map. The major capsid protein (MCP; ORF40) of the penton (PP) and the peripentonal hexons (P1-6), triplex proteins TRX1 (ORF20) and TRX2 (ORF41) for triplexes Ta and Tc, and small capsomere interacting protein (SCP; ORF23) are colored lavender blue (PP), very soft violet (P1,3,5), light coral (P2,4,6), deep sky blue (TRX1), Columbia blue (TRX2.1), aqua (TRX2.2), and dark gray (SCP). The map transitions from opaque to transparent revealing the atomic model. The majority of the map fades to leave behind a pseudo-asymmetric unit. A PP MCP, P2 MCP, and Ta triplex transition back to opaque then the pseudo-asymmetric unit rotates. The remaining transparent segmented proteins fade away, then the P2 MCP and Ta triplex fade away leaving behind the PP MCP. The PP MCP becomes transparent, rotates, the field of view zooms in, individual helices become opaque followed by their exploded views of the extracted densities. This repeats for the P2 MCP and the individual proteins for the Ta triplex. The field of view zooms out to the pseudo-asymmetric unit followed by the fade in of the remaining segmented cryo-ET map.

**Supplemental Movie 8. Molecular Dynamics Flexible Fitting of the CVSC.** The movie shows molecular dynamics flexible fitting of the AlphaFold predicted CVSC into a portion of the 8.3Å VZV capsid vertex cryo-ET map. The CVSC is composed of CVC1 (ORF43), CVC2 (ORF34; two copies), and VP1/1 (ORF22; two copies of the C-terminal alpha helix); CVC1 – orange, CVC1.1 – red, CVC2.2 – falu red, VP1/2.1 – yellow, VP1/2.2 – gold.
