## Supplementary Information for "Cryogenic Electron Tomography Redefines Herpesvirus Capsid Assembly Intermediates Inside the Cell Nucleus"

#### for

Oliver et al.

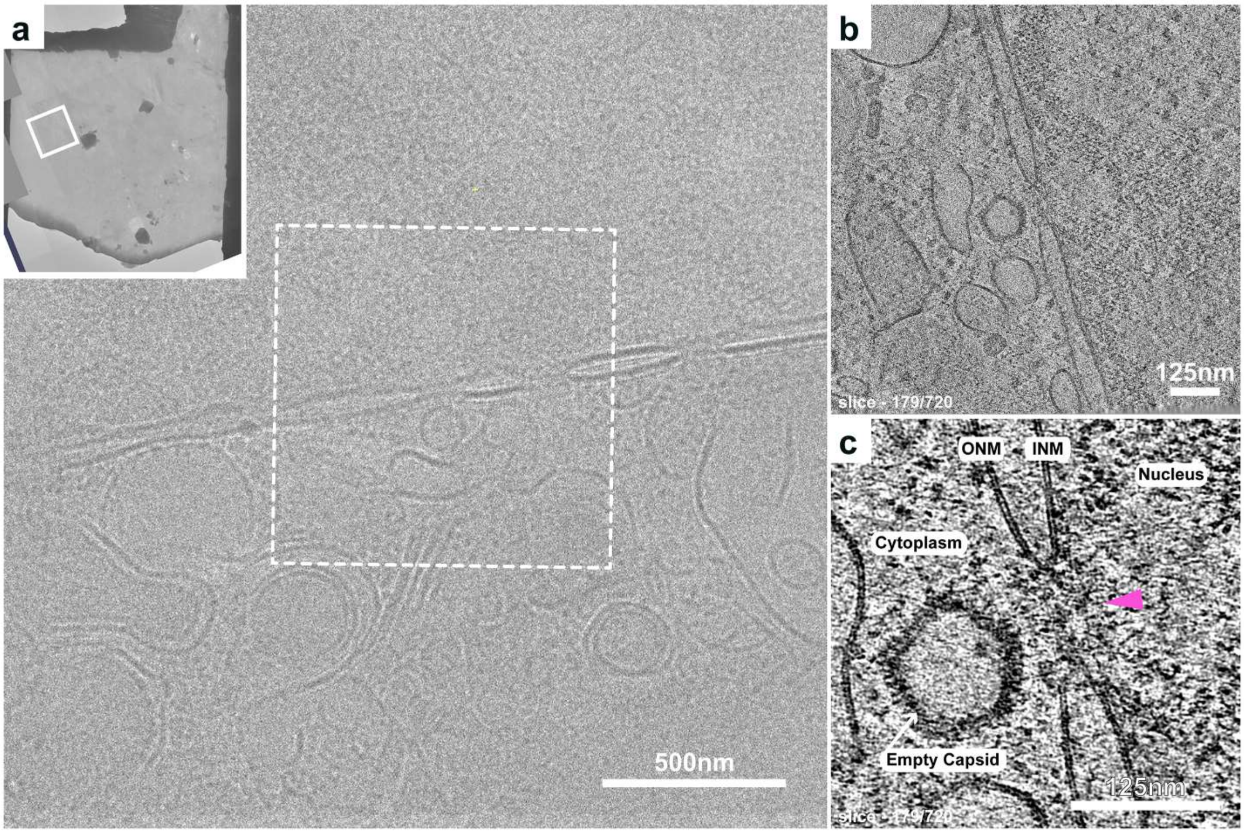

**Supplemental Figure 1. A VZV capsid docked at a nuclear pore visualized *in situ* by cryo-FIB-SEM/cryo-ET. (a)** A montage image (16.85Å/pixel) of a lamella produced by cryogenic transmission electron microscopy (cryo-TEM). The inset white box highlights the zoomed in area where an enveloped VZV capsid is visible at the nuclear periphery and cellular features in the cytoplasm. **(b)** A single slice (179/720) through a tomogram reconstruction of a tilt series captured from the dotted white box in panel A. An empty VZV capsid is visible at the nuclear pore. **(c)** A low-pass filtered image reveals the inner (INM) and outer (ONM) nuclear membrane converging at the nuclear pore (pink arrowhead) with the docked VZV capsid from panel B. Scale bars 500nm (**a**) and 125nm (**b and c**).

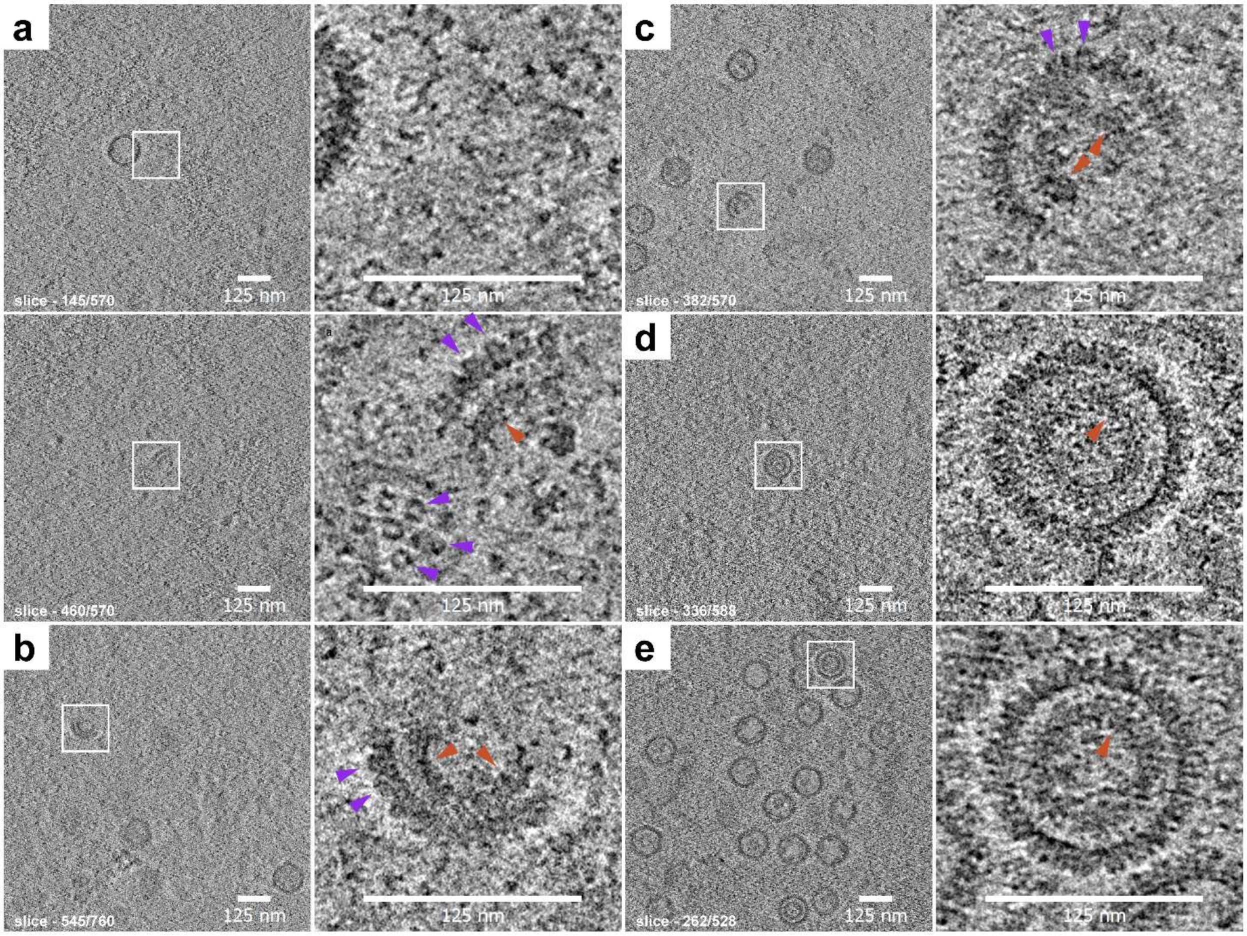

**Supplemental Figure 2. Intranuclear VZV procapsid intermediates identified by cryo-FIB/cryo-ET. (a to e)** Tomogram slices generated from cryo-ET of lamella produced using cryo-FIB/SEM of VZV infected MeWo cells. White boxes on the left-hand panels highlight the magnified images in the right-hand panels. The tomogram slice per total for each image is given in the bottom left corner of the panels. **(a to c)** Procapsid assembly precursors were identified in the nuclei of VZV infected cells. The capsomeres (blue violet arrowheads) and scaffold protein (Trinidad arrowheads) were visible on wedge (a) or half-moon shaped (b and c) macromolecular assemblies. To aid visualization two slices of the tomogram for A are shown revealing an A/B capsid next to the procapsid precursors; the white box is in the same location of the tomogram. **(d and e)** Examples of procapsids in the nuclei of VZV infected cells. Scaffold protein is highlighted by the blue arrowheads. All scale bars represent 125nm.

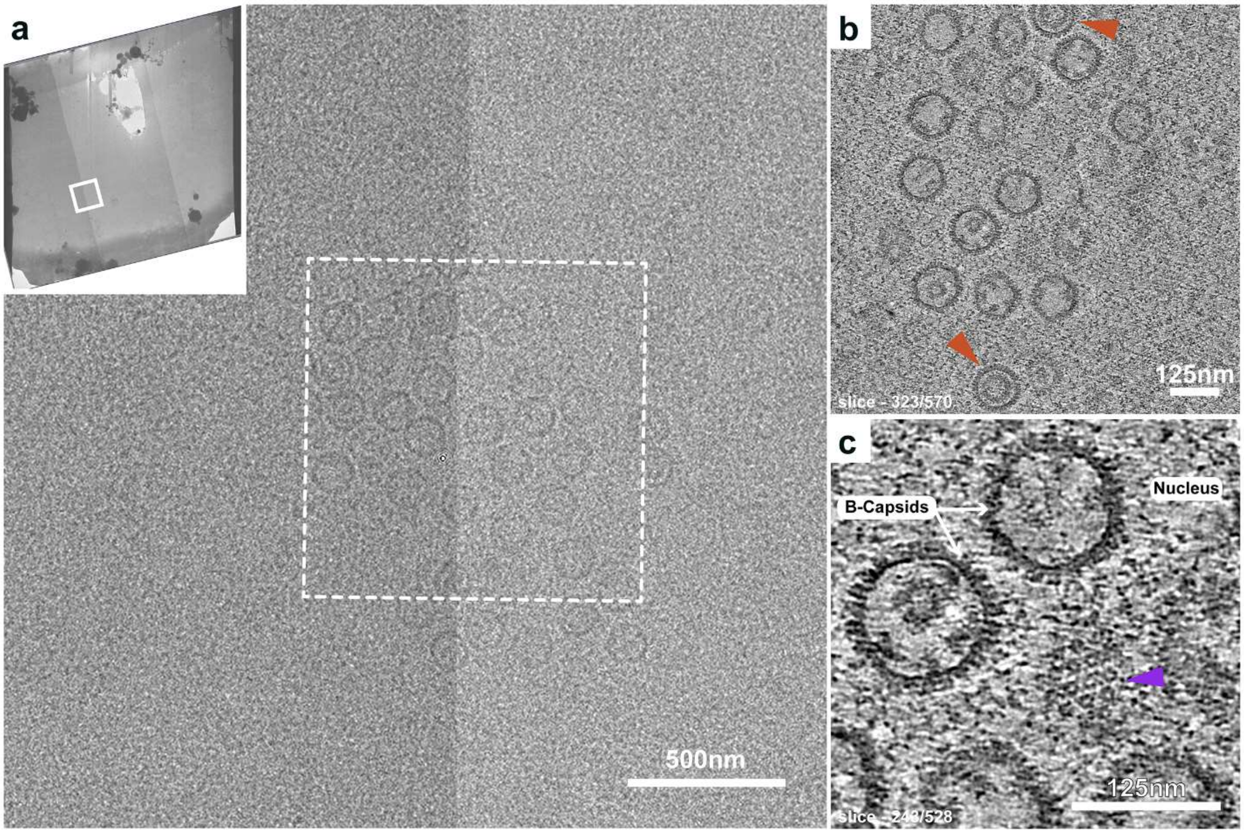

**Supplemental Figure 3. Newly assembled VZV capsids visualized *in situ* by cryo-FIB-SEM/cryo-ET. (a)** A montage image (39.14 Å/pixel) of a lamella produced by cryogenic transmission electron microscopy (cryo-TEM). The inset white box highlights the zoomed in area where an enveloped VZV capsid is visible at the nuclear periphery and cellular features in the cytoplasm. **(b)** A single slice (243/528) through a tomogram reconstruction of a tilt series captured from the dotted white box in panel A. A cluster of newly assembled VZV pro-capsids (Trinidad arrowheads) and B-capsids. **(c)** A low-pass filtered image highlights two B-capsids containing scaffold protein and the capsomeres of an adjacent capsid along the 3-fold axis (blue violet arrowhead). Scale bars 500nm (**a**) and 125nm (**b and c**).

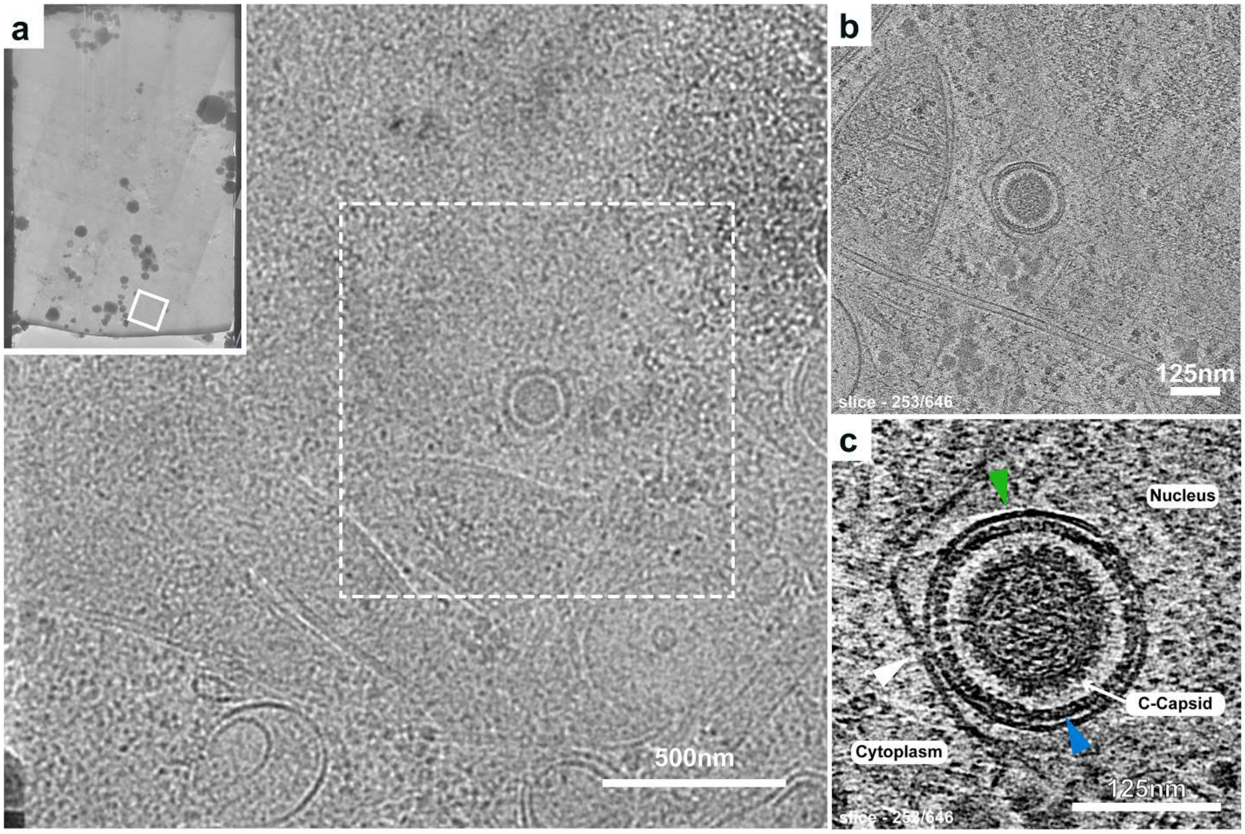

**Supplemental Figure 4. VZV nuclear egress visualized *in situ* by cryo-FIB-SEM/cryo-ET. (a)** A montage image (16.85Å/px) of a lamella produced by cryogenic transmission electron microscopy (cryo-TEM). The inset white box highlights the zoomed in area where an enveloped VZV capsid is visible at the nuclear periphery and cellular features in the cytoplasm. **(b)** A single slice (253/646) through a tomogram reconstruction of a tilt series captured from the dotted white box in panel A. A primary envelope around a VZV C-capsid is visible. **(c)** A low-pass filtered image reveals the VZV capsid from panel B surrounded by the nuclear egress complex (navy blue arrowhead) and the primary envelope (lime green arrowhead) undergoing fusion with the outer nuclear membrane (white arrowhead). Scale bars 500nm (**a**) and 125nm (**b and c**).

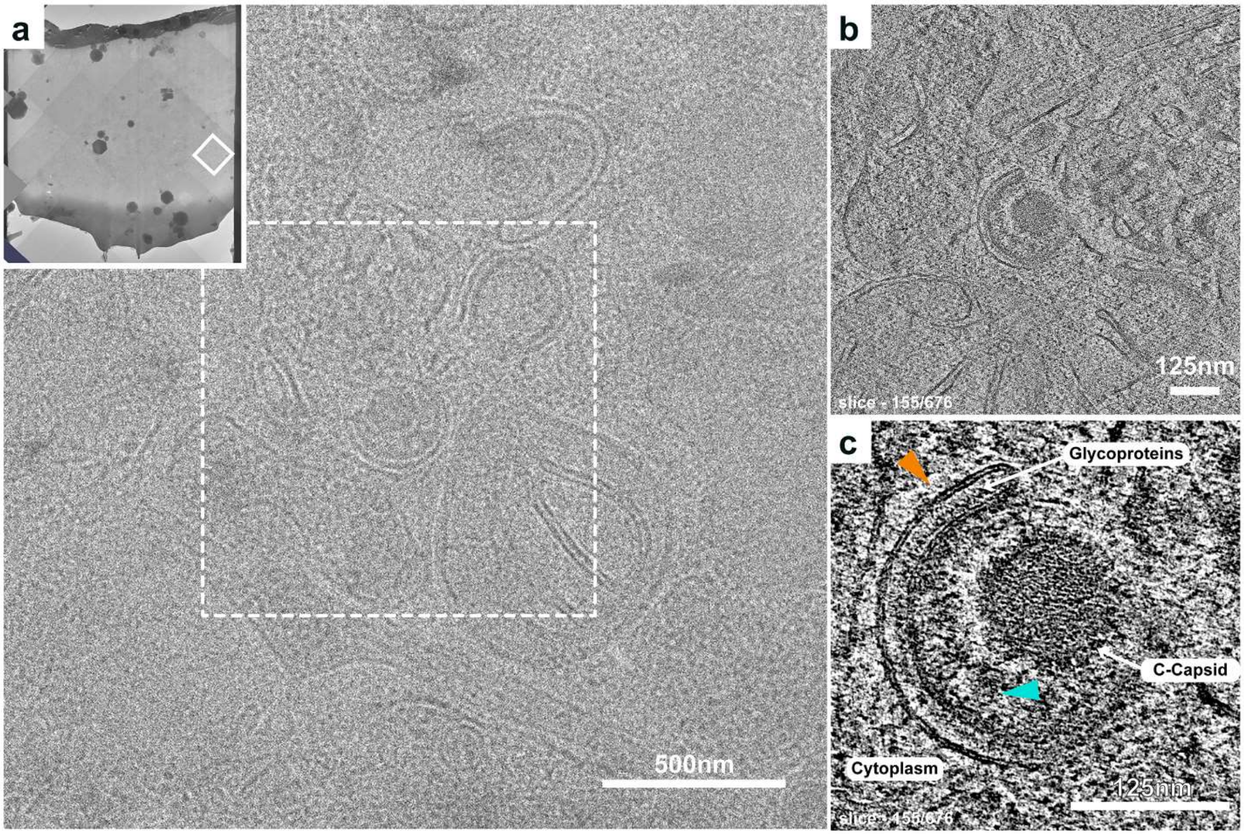

**Supplemental Figure 5. VZV secondary envelopment visualized *in situ* by cryo-FIB-SEM/cryo-ET. (a)** A montage image (16.846Å/px) of a lamella produced by cryogenic transmission electron microscopy (cryo-TEM). The inset white box highlights the zoomed in area where a VZV is in the process of secondary envelopment and cellular features in the cytoplasm. **(b)** A VZV C-capsid captured during secondary envelopment. A single slice (155/676) through a tomogram reconstruction of a tilt series captured from the dotted white box in panel A. **(c)** A low-pass filtered image reveals the VZV C-capsid interacting with the tegument layer (bright turquoise arrowhead) and glycoprotein-coated cell membranes (tangerine arrowhead) morphing around the capsid-tegument complex. Scale bars 500nm (**a**) and 125nm (**b and c**).

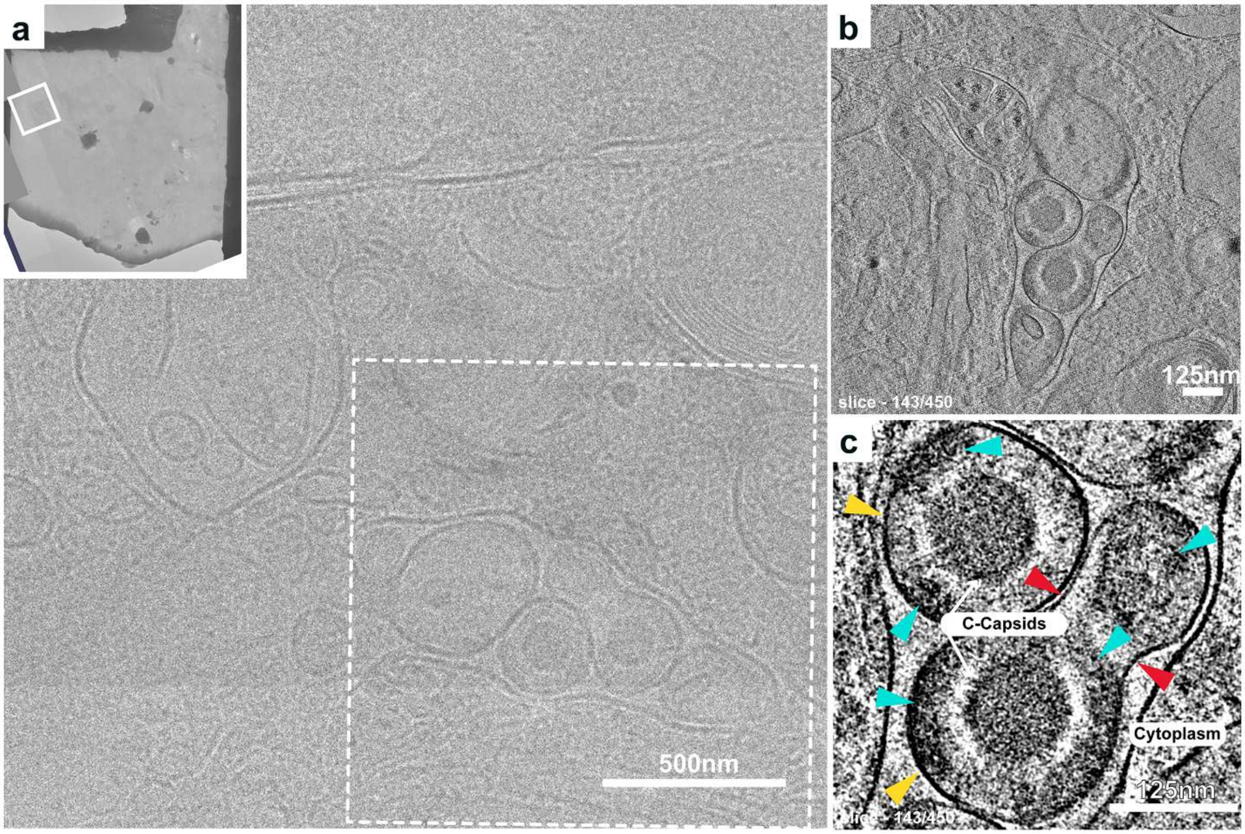

**Supplemental Figure 6. VZV morphogenesis visualized *in situ* by cryo-FIB-SEM/cryo-ET.** **(a)** A montage image (16.85Å/pixel) produced by cryogenic transmission electron microscopy (cryo-TEM). The inset white box highlights the zoomed in area where an enveloped VZV capsid is visible at the nuclear periphery and cellular features in the cytoplasm. The inset blue box highlights the zoomed in area where virus particles are captured in the process of morphogenesis. **(b)** VZV capsids captured in a vesicular structure during morphogenesis. A single slice (143/450) through a tomogram reconstruction of a tilt series captured from the dotted white box in panel A. **(c)** A low-pass filtered image of a VZV particle that has undergone complete morphogenesis and a second in the process of the final steps of morphogenesis and generation of a light particle, a VZV particle without a capsid. The tegument is visible as the dark structures (bright turquoise arrowheads) between the C-capsids and the VZV particle envelope (gorse yellow arrowheads). Red arrowheads indicate a point of constriction between the formation of a complete VZV particle and a light particle. Scale bars 500nm (**a**) and 125nm (**b and c**)

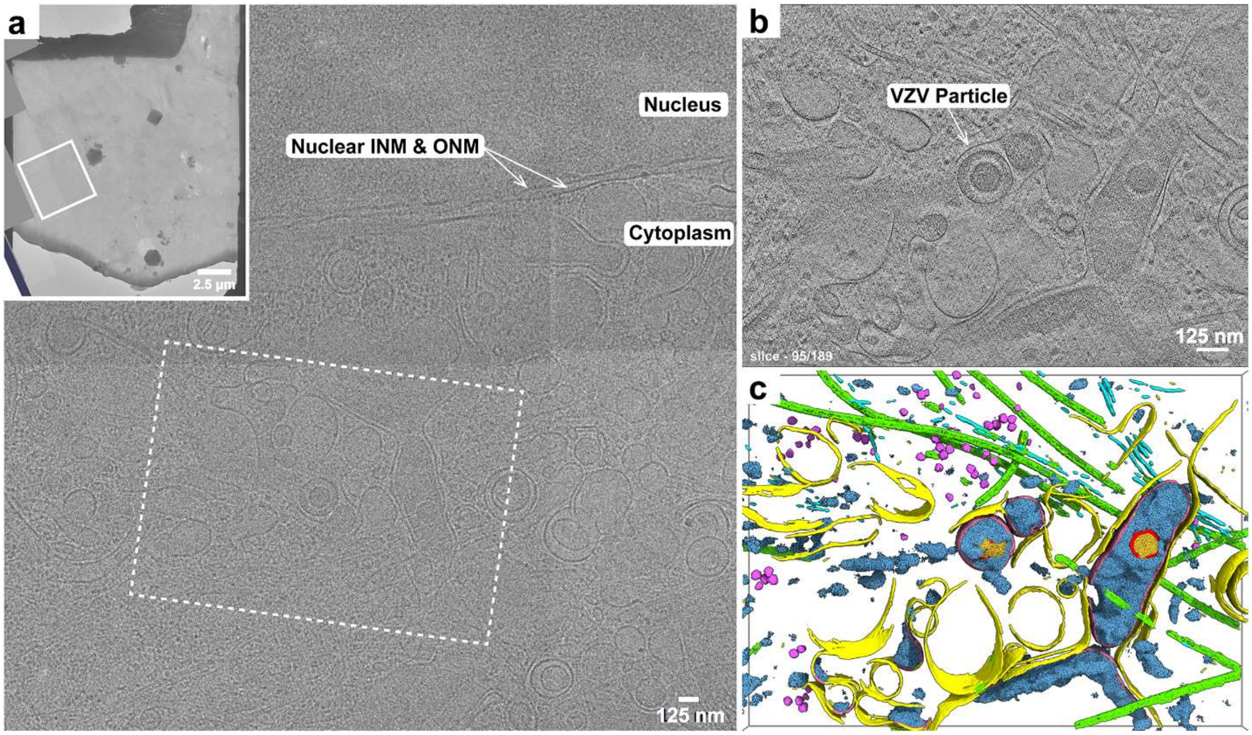

**Supplemental Figure 7. VZV morphogenesis visualized *in situ* by cryo-FIB/cryo-ET. (a)** A montage image (16.85Å/pixel) produced by cryo-TEM of the lamella (inset). The inset white box highlights the zoomed in area where the inner (IM) and outer (OM) nuclear membranes are visible and numerous cellular features are seen in the cytoplasm. Scale bar 125nm; inset 2.5µm. **(b)** A single slice (95/189) through a tomogram reconstruction of a tilt series captured from the dotted white box in panel **a**. A complete VZV particle is indicated in the tomogram. Scale bar 125nm. **(c)** Segmentation (Dragonfly) of the tomogram in panel D. Eight features were segmented in the tomogram; microtubules (green), filaments (cyan), vesicle membranes (yellow), ribosomes (medium orchid), VZV envelope (tapestry), VZV tegument (picton blue), VZV capsid (red), and VZV DNA (bright sun).

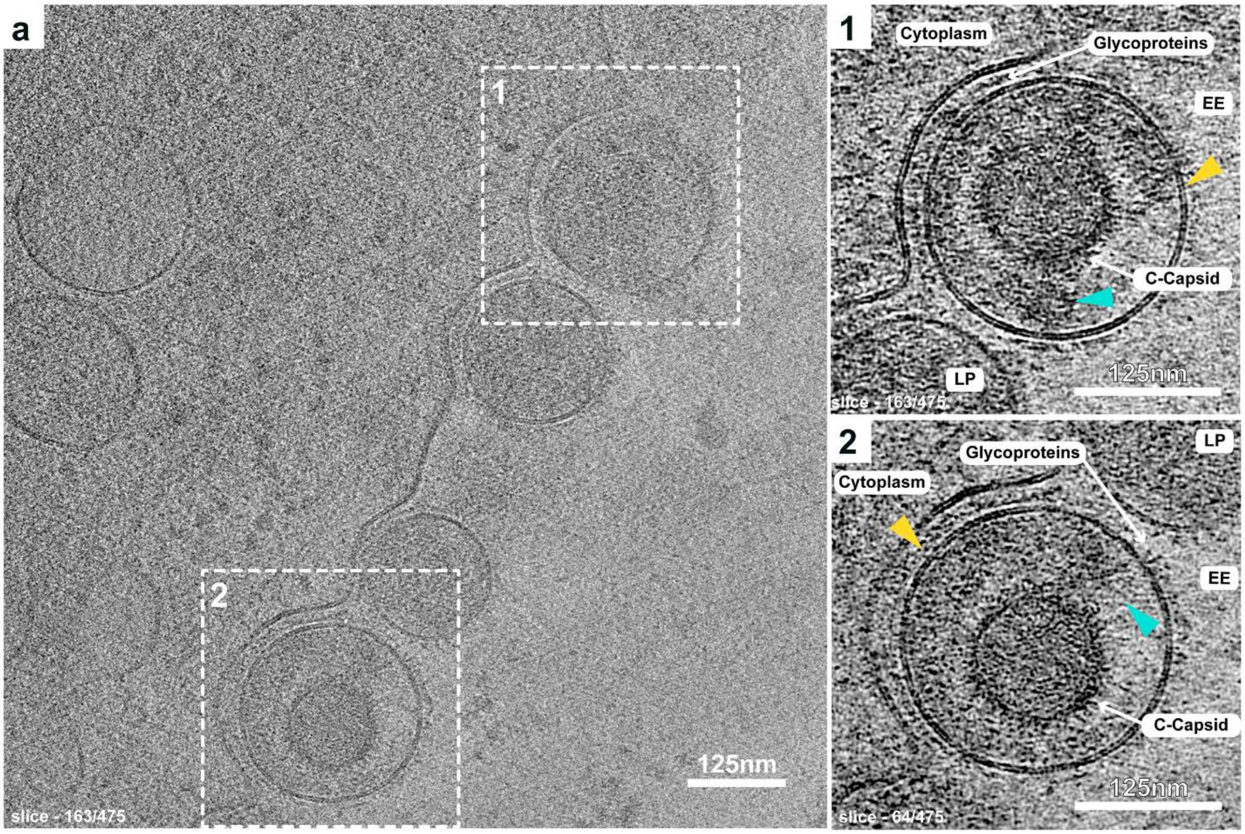

**Supplemental Figure 8. VZV particles at the plasma membrane visualized *in situ* by cryo-ET. (a)** A single slice (64/475) through a tomogram reconstruction of a tilt series generated by cryo-ET at the periphery of a vitrified VZV infected MeWo cell. VZV particles containing C-capsids released into the extracellular environment (EE) bound to the plasma membrane, which is typical for this cell associated herpesvirus. The dashed boxes (1 and 2) highlight the area of the low-pass filtered images in panels 1 (slice 64/475) and 2 (slice 163/475), revealing two plasma membrane bound extracellular VZV particles. The tegument is clearly visible as the dark structures between the capsids (bright turquoise arrowheads) and the VZV particle envelope (gorse yellow arrowheads), which is coated with glycoproteins. Light particles (LP) also bound to the plasma membrane are partially visible in both panels. Scale bars 125nm (**a**) and (**1 and 2**).

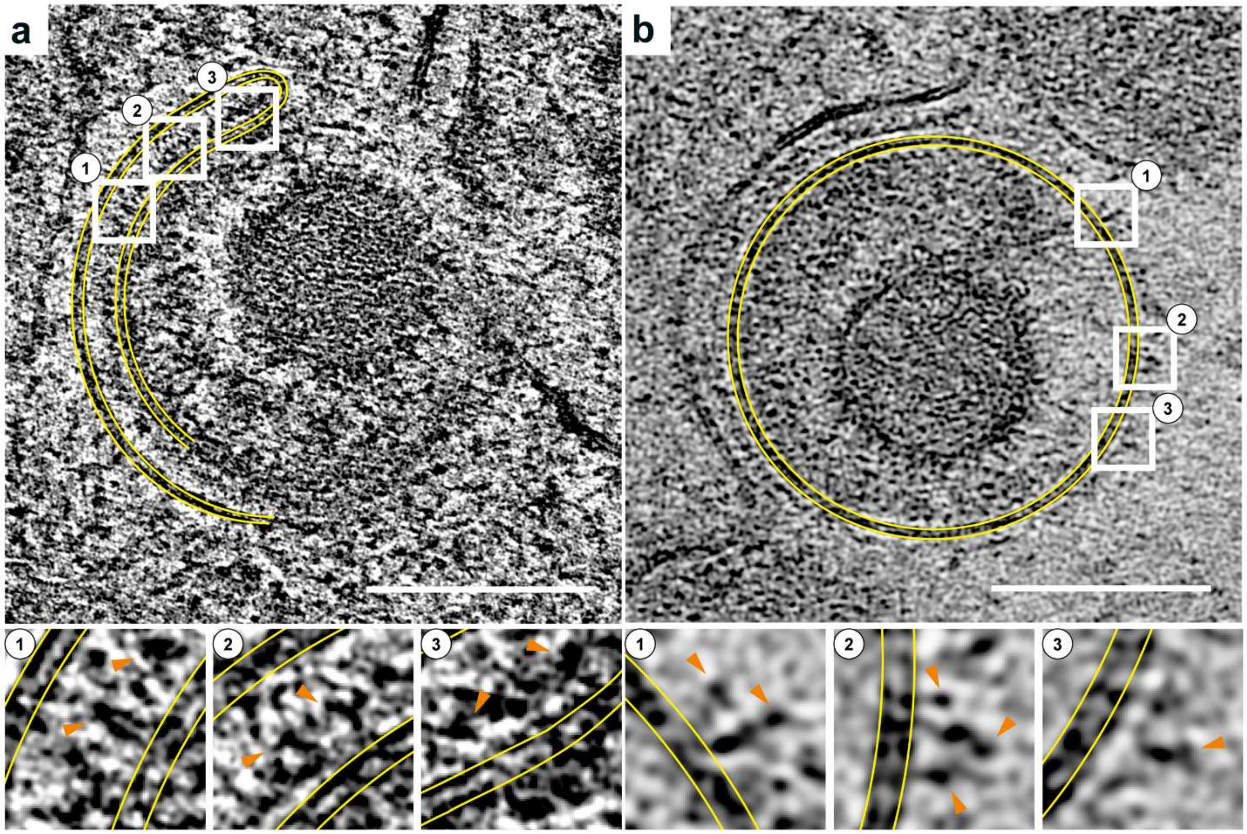

**Supplemental Figure 9. Glycoproteins on VZV envelopes.** Glycoproteins were visible the envelopes of VZV particles during secondary envelopment (**a**) and nascent cell associated particles (**b**). The yellow lines outline the lipid bilayers. Boxes 1 to 3 for each panel provide magnified views to visualize individual glycoprotein. The tangerine arrow heads point to representative glycoproteins protruding from the lipid bilayer for the envelope of a VZV particle undergoing secondary envelopment (**a**) or the fully formed VZV particle in the extracellular space. (**b**).

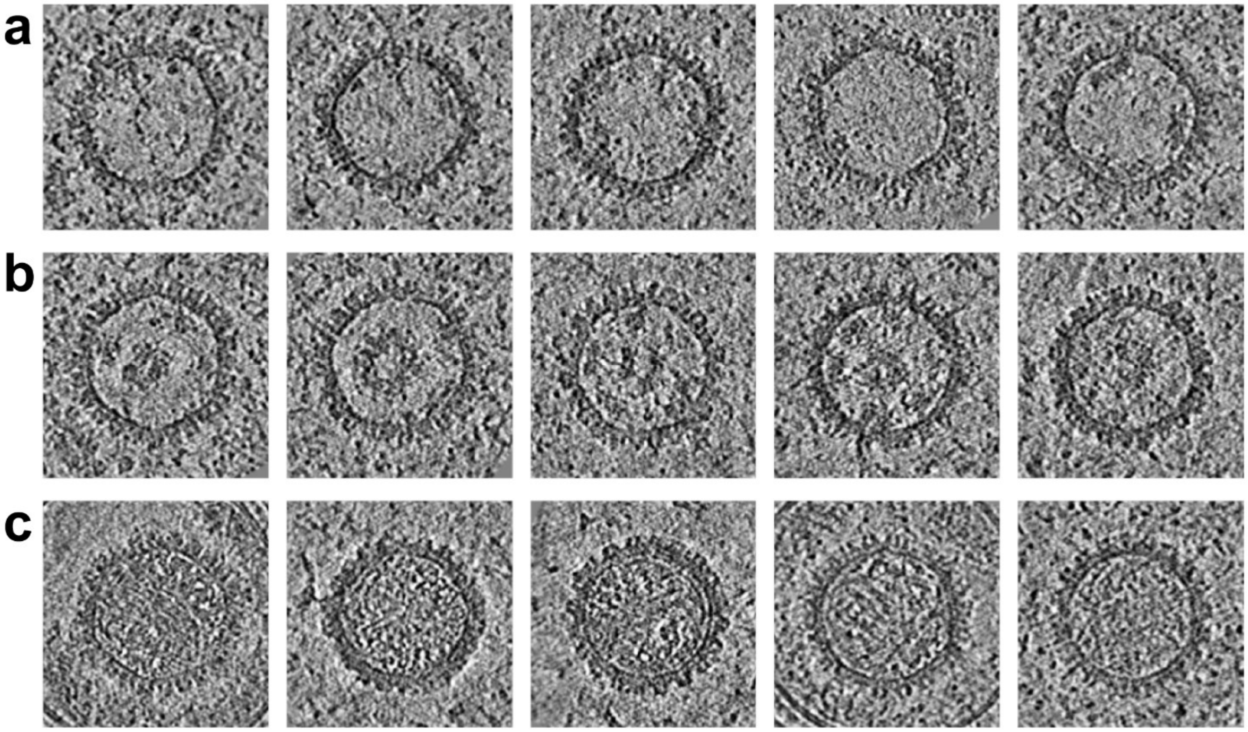

**Supplemental Figure 10. Examples of A-, B-, and C-like capsids.** The images were derived from tomograms generated in EMAN2 and shows a single slice through the center of the subtomograms for individual capsids. Each row shows five representative capsids as defined by their classical morphology as either A-, B-, or C-capsids based on the absence (**a**) or presence (**b**) of scaffold protein or dsDNA genome (**c**) in the capsid cavity.

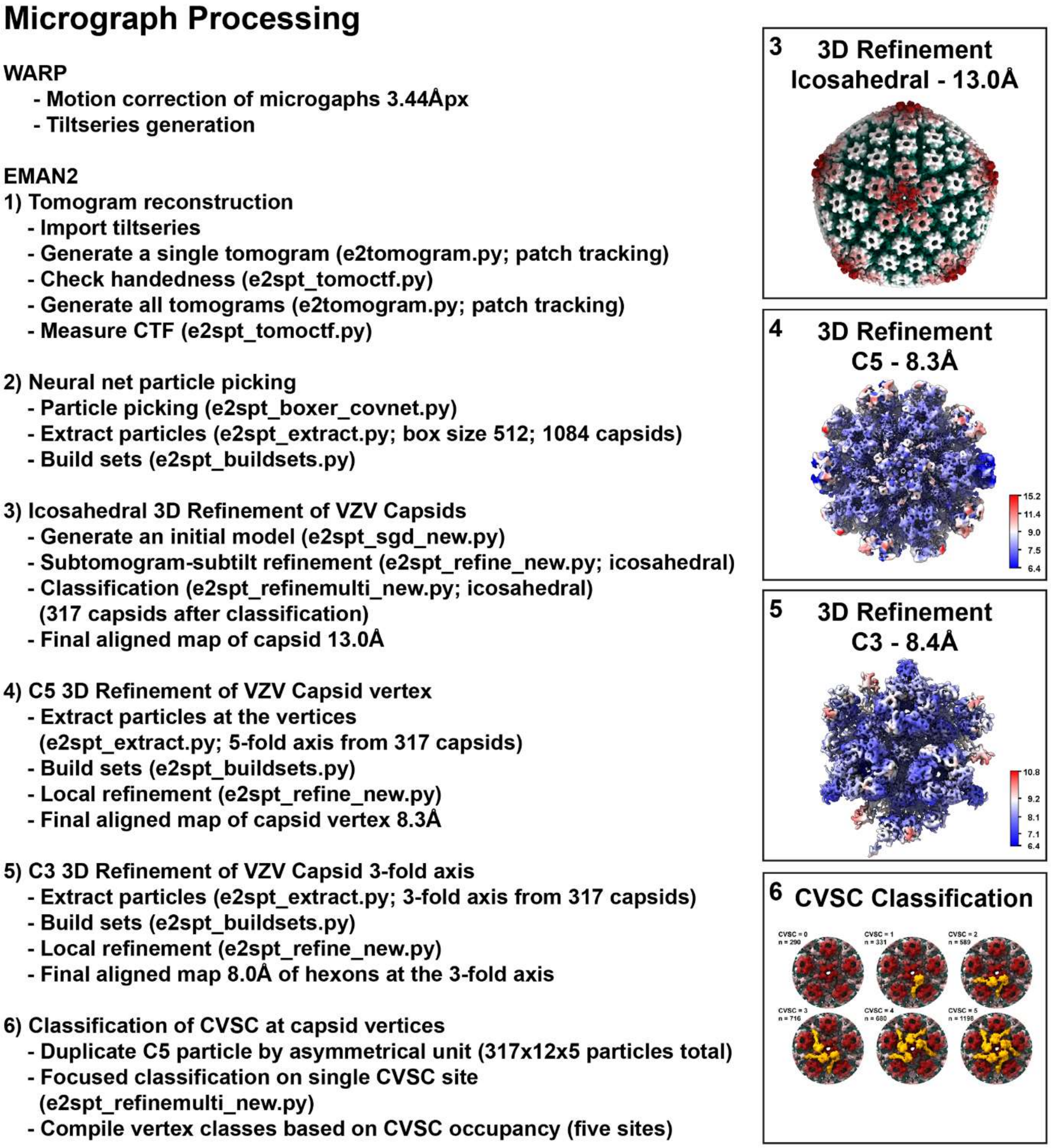

**Supplemental Figure 11. EMAN2 processing scheme for the reconstruction of cryo-ET maps of the VZV capsid (13Å), capsid vertex (8.3Å), and the 3-fold (8.4Å) symmetry axis.**

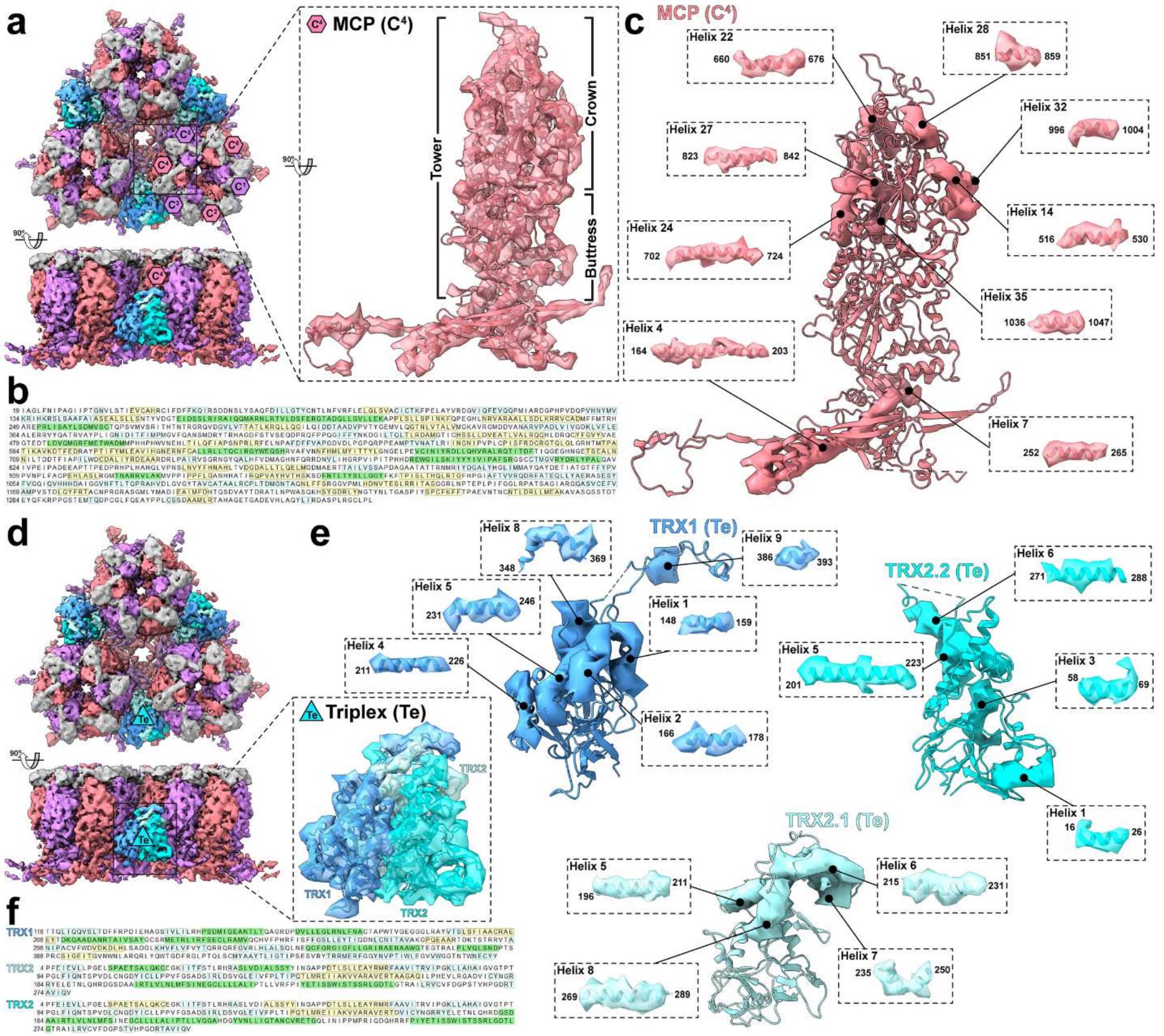

**Supplemental Figure 12. Expected features for the hexons and triplexes revealed by segmentation of the subnanometer 3-fold symmetry axis cryo-ET map. (a to f)** Segmentation of the 8.0Å cryo-EM map based on the 7BW6[1] and a refined model based on the subnanometer map. **(a and d)** The major capsid protein (MCP) of the central hexons ($C^{1-6}$), triplex proteins TRX1 and TRX2 (Te), and small capsomere interacting protein (SCP) are colored very soft violet ($C^{1,3,5}$), light coral ($C^{2,4,6}$), deep sky blue (TRX1), Columbia blue (TRX2.1), aqua (TRX2.2), and dark gray (SCP). **(b and f)** Amino acid sequences for the central hexon MCP ($C^4$) and triplex Te (TRX1, TRX2.1, and TRX2.2) with segmented helices highlighted in green. **(e and e)** Exploded views of the extracted densities for helices from the MCP ($C^4$) and triplex Te subunits. Each helix is numbered according to the primary (**d and f**) structure along with the beginning and end numbered amino acids.

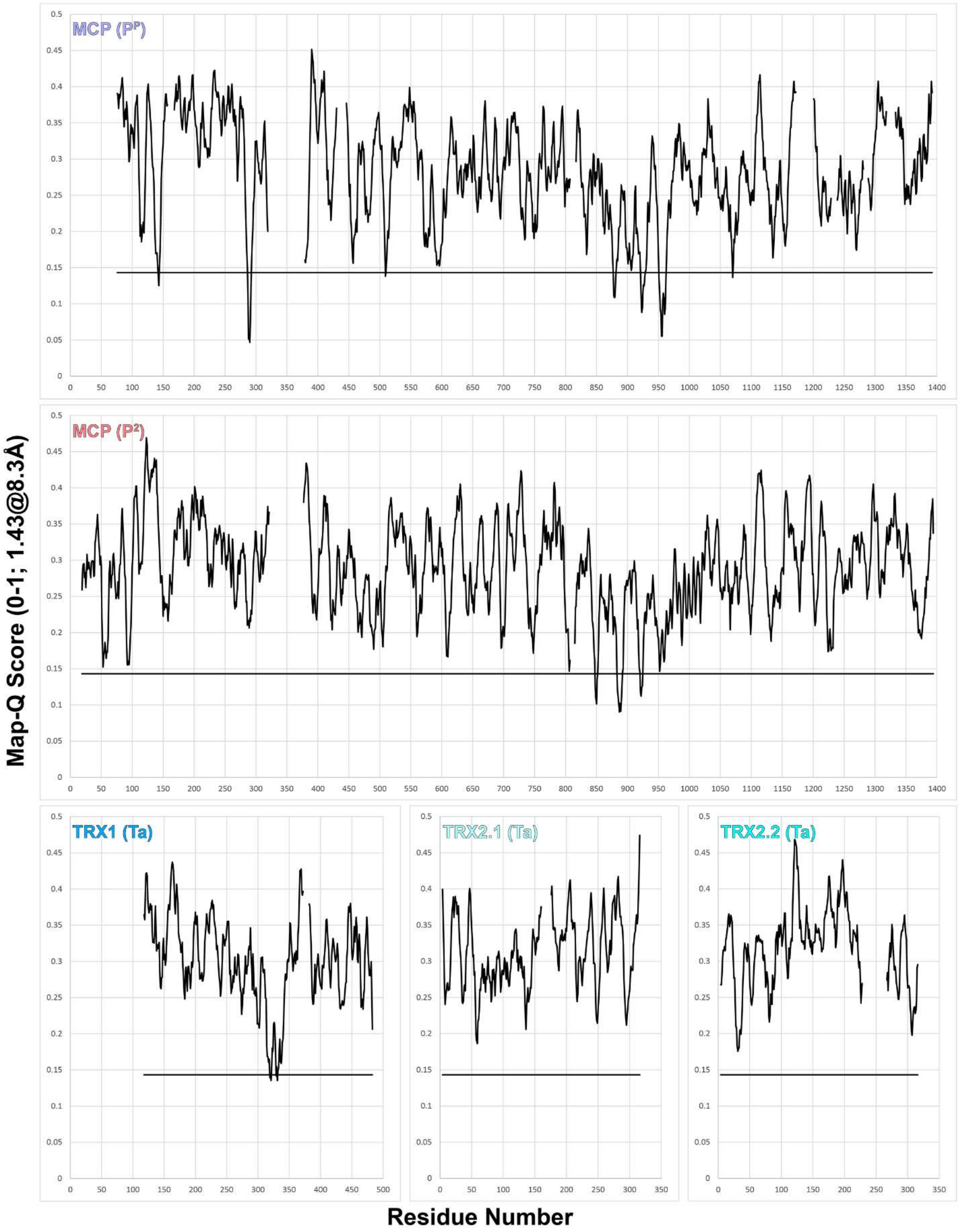

**Supplemental Figure 13. Map-Q scores for the pentonal and peripentonal MCP and triplex proteins**. Map-Q scores generated for the pentonal ($P^P$) and peripentonal ($P^2$) MCPs, and Ta triplex proteins TRX1, TRX2.1, and TRX2.2 were calculated using the indicated protein and the 8.3Å vertex cryo-ET map of the VZV capsid (**Fig 3 and 4**). The horizontal black line at 0.145 is the expected value of the Map-Q score for a structure at 8.3Å resolution. Gaps in the traces indicate regions in the protein structure models where amino acids were absent.

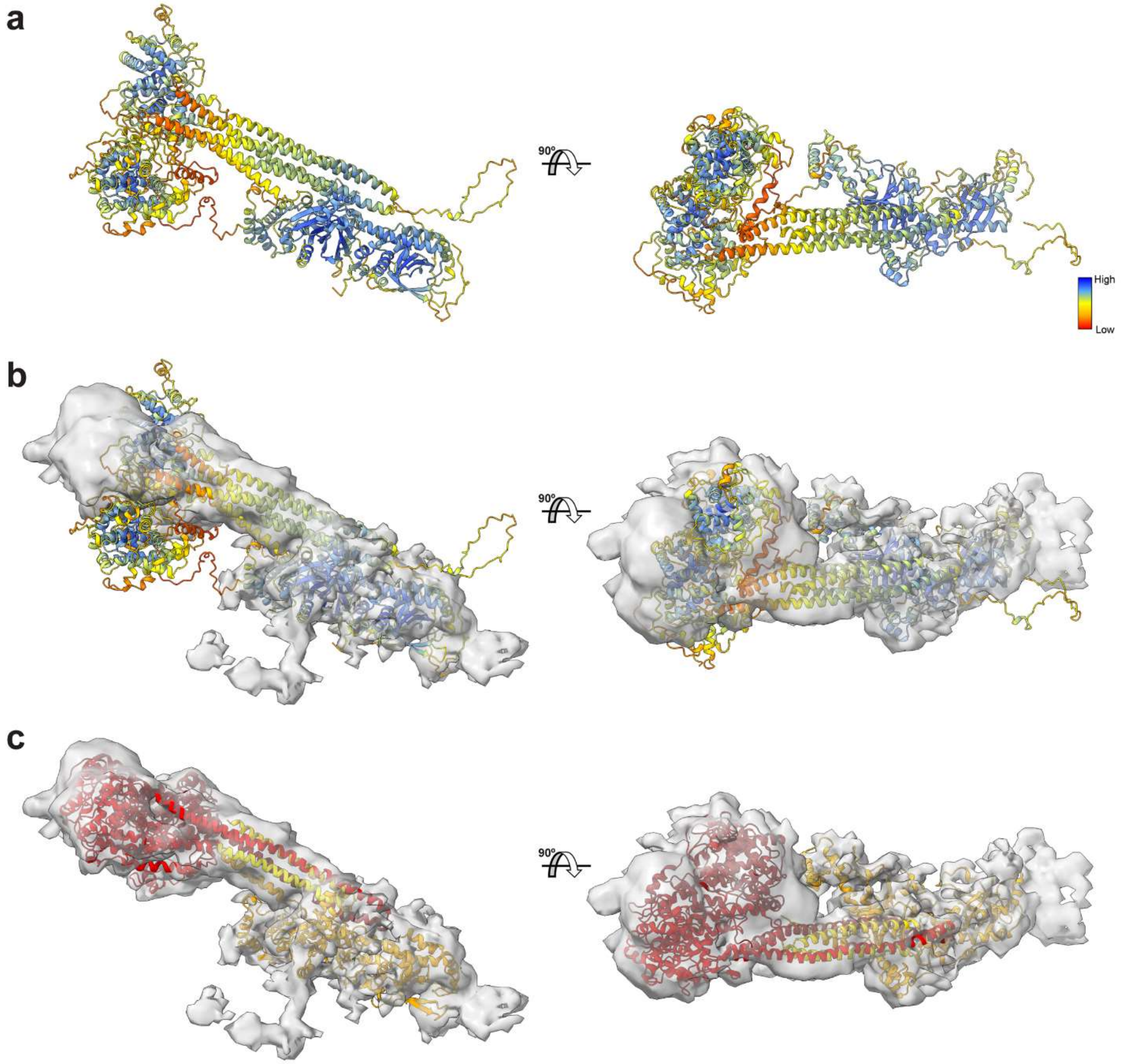

**Supplemental Figure 14 AlphaFold prediction and molecular dynamics flexible fitting of the VZV CVSC into the vertex cryo-ET map.** **(a)** AlphaFold-based prediction of the CVSC performed in multimer mode with the CVC1 (ORF43), CVC2 (ORF34; two copies), and VP1/1 (ORF22; two copies of the C-terminal alpha helix). **(b)** Fitment of the AlphaFold 2 two model (A) into the CVSC density segmented form the 8.3Å vertex cryo-ET map. A and B – Color code represents AlphaFold confidence levels. **(c)** Fitment of the CVSC after MDFF performed with NAMD. CVC1 – orange, CVC1.1 – red, CVC2.2 – falu red, VP1/2.1 – yellow, VP1/2.2 – gold.

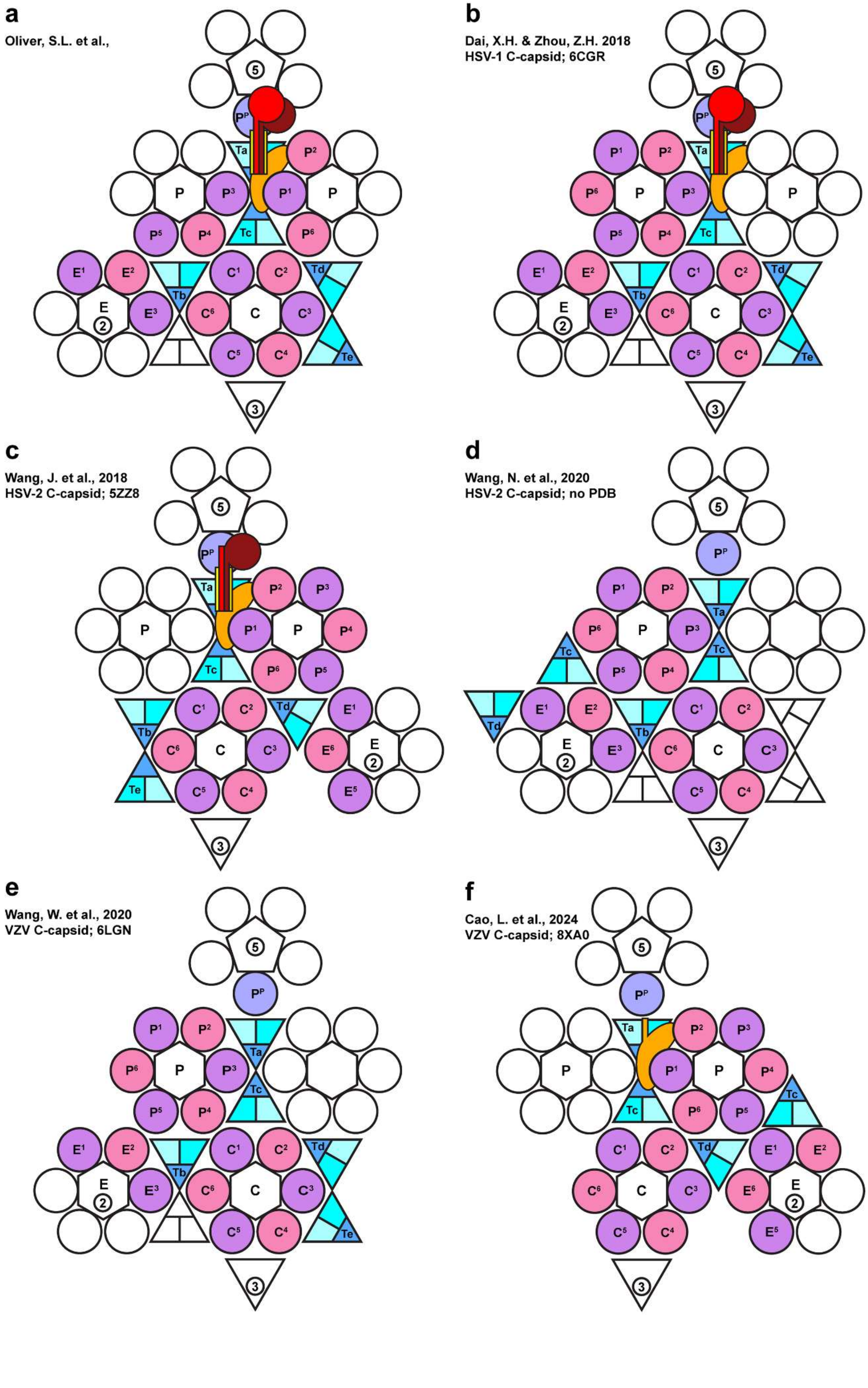
a
Oliver, S.L. et al.,
b
Dai, X.H. & Zhou, Z.H. 2018
HSV-1 C-capsid; 6CGR
c
Wang, J. et al., 2018
HSV-2 C-capsid; 5ZZ8
d
Wang, N. et al., 2020
HSV-2 C-capsid; no PDB
e
Wang, W. et al., 2020
VZV C-capsid; 6LGN
f
Cao, L. et al., 2024
VZV C-capsid; 8XA0

**Supplemental Figure 15. Cartoon representations of human alphaherpesvirus asymmetric units. a to f** – Asymmetric units derived for human alphaherpesviruses by cryo-ET in the current study (a) or cryo-EM in previous studies (b to f) are represented by colored shapes that define the pentonal major capsid protein ($P^P$), the three types of hexon (peripentonal ($P^1$-$P^6$), edge ($E^1$-$E^3$), and center ($C^1$-$C^6$)), the five triplexes (Ta, Tb, Tc, Td, and Te), and the CVSC subunits CVC1 (orange) CVC2.1 (red), CVC2.2 (falu red), VP1/2.1 (yellow), and VP1/2.2 (gold).

### Micrograph Processing

**WARP**
- Motion correction of microgaphs 3.44Åpx
- Tilt series generation

**IMOD**
- Tilt series alignment
- Ctf Plotter

**Prepare STAR files**
- Tomogram STAR file

**RELION**
- Import tomograms
- Tomogram reconstruction (bin 4)

**EMAN2**
- Import tomograms
- Particle picking (box size 120; 471 capsids)

**RELION**

**1) De Novo Model Generation; capsid**
- Import tomograms (tomogram STAR file)
- Import particles (particles STAR file)
- Generate pseudotomograms (bin 4: 13.76Åpx)
- 3D initial model (I3)
- 3D auto-refine (I3; mask)

**2) Icosahedral 3D Refinement; capsid**
- 3D auto-refine (I3)
- Reconstruct Particle
- Generate pseudotomograms
- 3D auto-refine (I3; mask)
- Tomo frame alignment

**3) C5 3D Refinement; capsid vertex**
- Symmetry expansion I3
- Select unique vertices (5,318)
- Adjust Z (3dmod to define pixels)
- Reconstruct Particle
- Generate pseudotomograms
- 3D auto-refine (C5, mask)

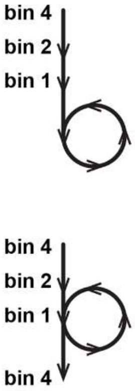

**4) 3D Classification; capsid vertex**
- 3D classification (12 classes, 29 iterations)
- Select particles (Class III & XI; 103 vertices)
- Template based 3D Classification
- Remove duplicate particles

**5) Portal Vertex 3D Refinement (109 vertices)**
- Reconstruct Particle
- Generate pseudotomograms
- 3D auto-refine (C5, mask)
- Final aligned map of portal vertex 23.6Å

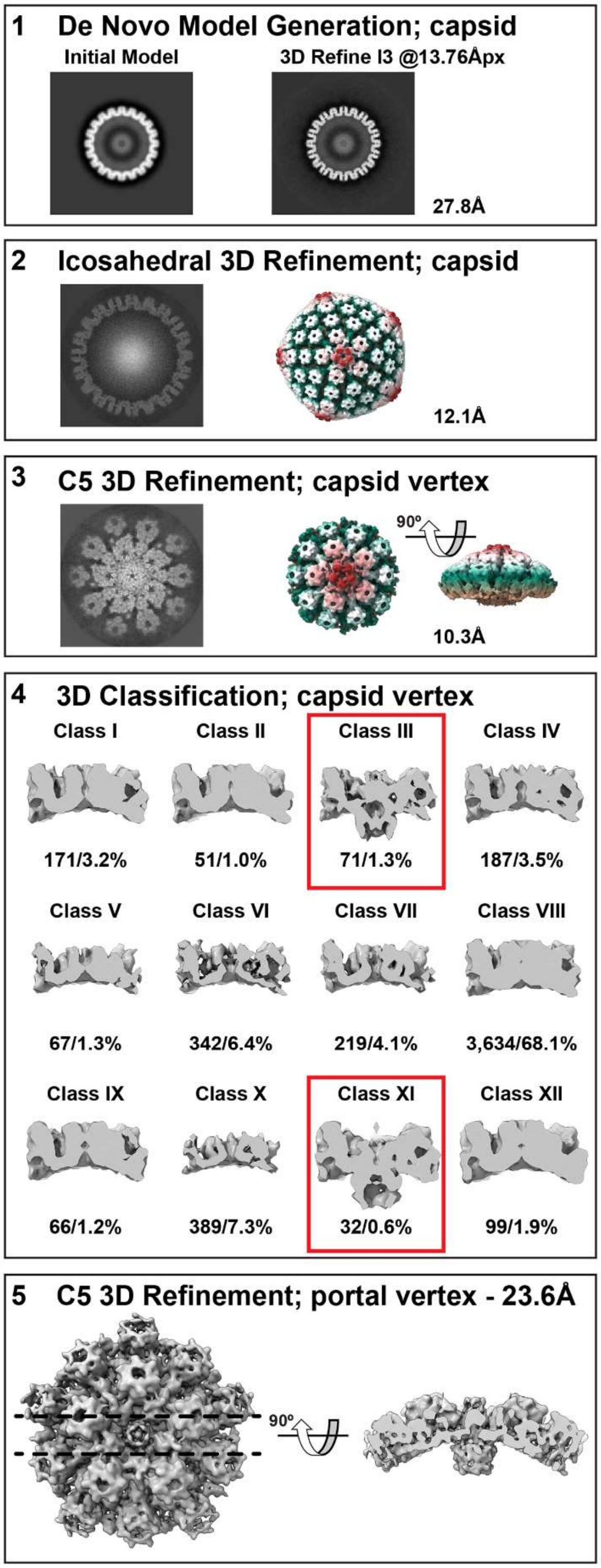

**Supplemental Figure 16. Relion processing scheme for the reconstruction of the VZV portal vertex cryo-ET map.** An outline of the Relion-based micrograph processing protocol used for the reconstruction of the VZV portal vertex from intracellular capsids revealed by cryo-FIB/cryo-ET of VZV infected cells. Tilt series from motion corrected movie data captured by a Titan Krios (300kV) electron microscope with a 20eV energy filter and a dose symmetric tilt series of 2°, ±60° were generated by WARP. Tilt series (n=94) were aligned with IMOD and tomograms (bin 4) from the aligned tilt series were generated by Relion. B- and C-capsids (n=471) were picked from the tomograms using EMAN2 and the coordinates imported into Relion for refinement: 1 – De novo model generation was performed from pseudo-subtomograms (bin 4; 13.76Å) with the 471 capsids using icosahedral symmetry (I3) and reached resolution near Nyquist (27.8Å); 2 – Several rounds of icosahedral 3D refinement was performed to generate a cryo-ET map of the VZV capsid at 12.1Å (EMD-49465); 3 – Symmetry expansion and removal of vertices that were beyond the tomogram boundaries produced 5,318 unique vertices that were used for C5 symmetry 3D refinement (EMD-49466). Several rounds of refinement were performed to generate a cryo-ET map of the VZV capsid vertex at 10.3Å; 4 – 3D classification of the aligned vertices (bin 4) into 12 classes using 29 iterations produced two classes (n=103) with portal-like densities. Further template-based classification yielded a total of 109 portal vertices; 5 – A 23.6Å cryo-ET map of the VZV portal vertex was generated from the 109 capsids, revealing the portal density.

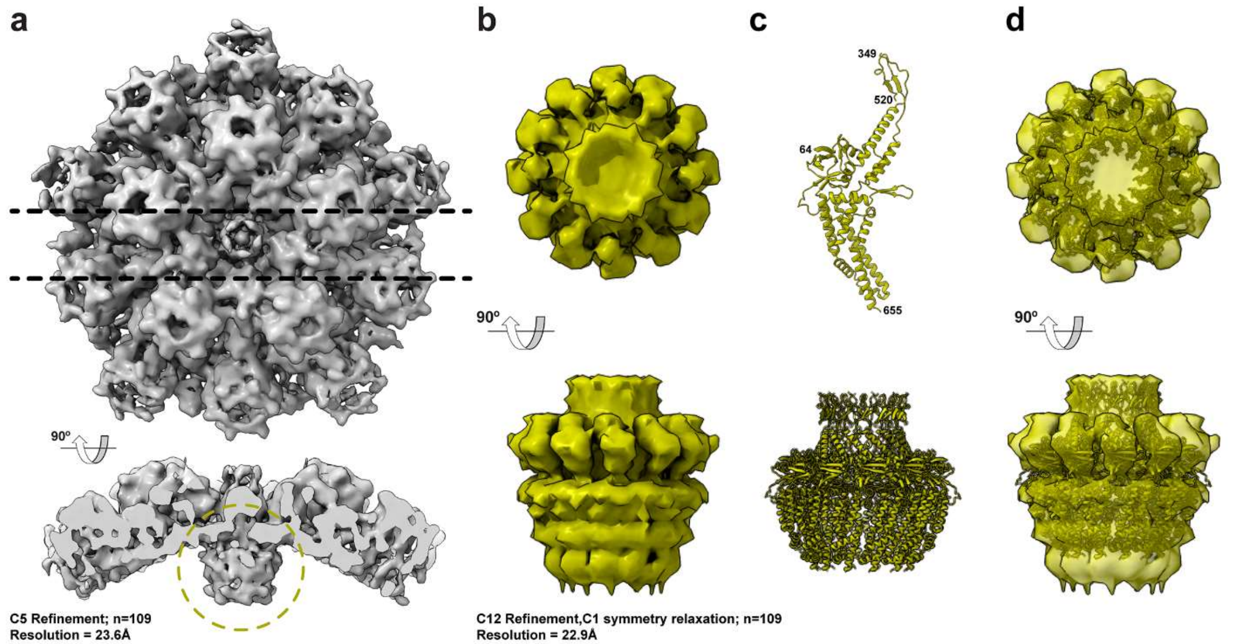

**Supplemental Figure 17. Cryo-ET reconstruction of the VZV portal vertex.** **(a)** – A 23.6Å cryo-ET map (EMD-49467) of the VZV portal vertex reconstructed from 109 capsids (CAIs n=62; C-capsids n=47) using the Relion-based processing pipeline outlined in Suppl. Fig. 14. The portal vertex is viewed along the Z-axis (upper panel) and the Y-axis (lower panel). The dashed lines in the upper panel show the slices used in the lower panel to more clearly visualize the portal highlighted by the dashed olive/yellow circle. **(b)** – A 22.9Å cryo-ET map (EMD-49468) of the VZV portal viewed along the Z-axis (upper panel) and the Y-axis (lower panel). The portal was reconstructed using 3D refinement with C12 symmetry and C1 symmetry relaxation. **(c)** – An AlphaFold-based prediction of the VZV portal protein (ORF54) and a molecular model of the VZV portal. ORF54 is 769aa in length. Disordered regions of the predicted structure were removed and the numbers indicate the amino acid residues (64-349 and 520-655) that make up the molecular model. The molecular model of the portal (lower panel) was based on the alignment of the VZV OR54 monomer model with the HSV-1 dodecamer of the pUL6 portal protein (PDB 6OD7). **(d)** – The VZV portal molecular model (panel **c**) fit into the cryo-ET map of the VZV portal (panel **d**).

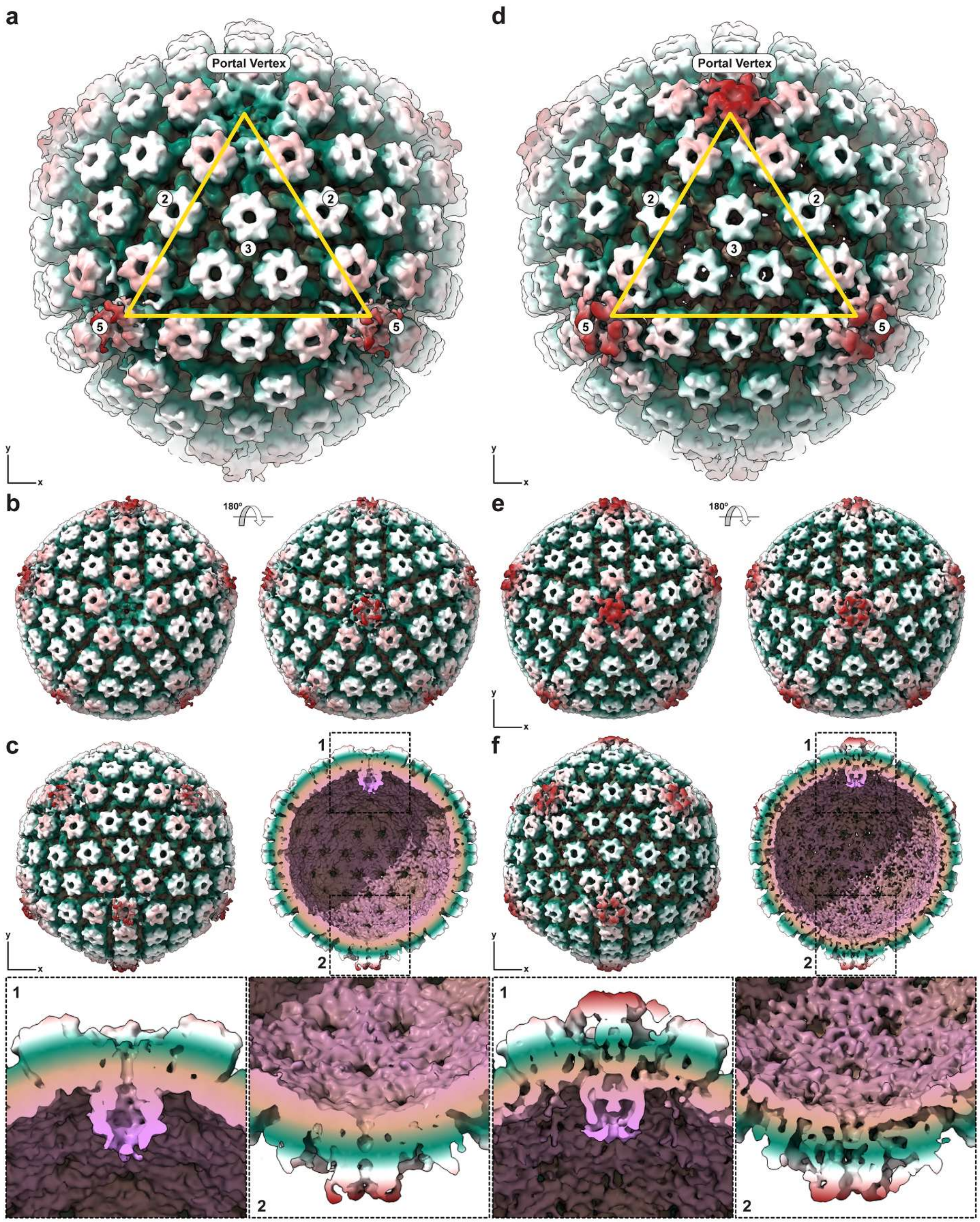

**Supplemental Figure 18. Reconstruction of cryo-ET maps for VZV CAIs and C-capsids revealing the unique portal vertex. (a to f)** Cryo-ET maps generated from VZV CAIs (n=62; 32.7 Å; EMD-49470) and C-capsids (n=47; 33.6Å; EMD-49472) reconstructed using 3D refinement with C5 symmetry and C1 symmetry relaxation. The cryo-ET maps are colored radially from the center of the capsid to the outer shell; pink, beige, green, white, and red. **(a and d)** The cryo-ET maps viewed along the 3-fold axis of symmetry with the portal vertex and the 2-, 3-, and 5-fold axes of symmetry highlighted by the numbered circles. **(b and e)** Capsids viewed along the Z-plane at the 5-fold axis of symmetry, which orientates the portal vertex along the Z-plane facing forwards (right-hand panels) and then rotated 180° (right-hand panels). **(c and f)** The portal vertex of the CAIs and C-capsids were aligned to the Z-plane then rotated 90°, which orientates the portal vertex along the Y-plane (upper-panels). The left-hand panels show the entire capsids and the right-hand panels show a slice view through the Y-plane. The two numbered boxes outline the portal vertex (1) and the polar opposite vertex (2) with high magnification views in the lower panels. The dsDNA genome is not visible inside the C-capsids due to the threshold parameters used to visualize the cryo-ET map.

**Supplemental Table 1. Cryo-ET data collection parameters.**

| Parameter | Microscope | | |
|---|---|---|---|
| | Talos Arctica[A] (FEI; TEM3) | Titan Krios (FEI; TEM2) | Titan Krios (FEI; TEMBETA) |
| Voltage (kV) | 200 | 300 | 300 |
| Magnification | 39,000 | 26,000 | 26,000 |
| Gatan Detector (pixels) | K2 Summit (3708 3838) | K3 (5760 4092) | K3 (5760 4092) |
| Energy filter slit width (eV) | 20 | 20 | 15 |
| Pixel size (Å) | 3.5 | 3.44 | 3.44 |
| Defocus range (µm)[B] | -4 | -3 to -5 | -2 to -3 |
| Exposure time (s) | 3 | 0.700 to 0.999 | 1.298 |
| Frames | 12 | 6-10 | 13 |
| Dose Rate $e^-$/pixel/s | 5.278 | 14.669 to 26.776 | 14.496 |
| Electron dose ($e^-$/Å$^2$) | 1.293 | 1.231 to 1.772 | 1.590 |
| Electron exposure rate ($e^-$/Å$^2$/s) | 0.431 | 1.240 to 2.263 | 1.225 |
| Dose per frame ($e^-$/Å$^2$/frame) | 0.108 | 0.119 to 0.271 | 0.122 |
| Tilt angle (°±) | 0-60 | 0-60 | 0-60 |
| Tilt step (°)[C] | 2 | 2 | 2 |
| Total electron exposure ($e^-$/Å$^2$)[D] | 1.293 | 0.867 to 1.927 | 1.590 |
| Total electron exposure ($e^-$/Å$^2$)[E] | 78.847 | 52.907 to 104.267 | 96.992 |

[A] Used for cryogenic electron tomography (cryo-ET) of cells without cryo-FIB-milling
[B] Target defocus
[C] Dose symmetric
[D] Per tilt
[E] Total dose per tilt series

**Supplemental Table 2. Tomograms used to represent steps in VZV particle morphogenesis.**

| Tilt series/Tomogram[A] | Movie[B] | Slice[C] | Image Processing [D] | Description |
|---|---|---|---|---|
| 20220507_CS02_VZV52G2_lamella3_tilt02 (16.846 Å/px) | 130-290 | 179/720 | 0.00/0.075/0.05 | A-capsid at a nuclear pore, cytoplasmic side. |
| 20211210_CS02_VZV46G2_Tilt19 Lamella: 20211210_CS02_Pos3_MMM3 (39.140 Å/px) | 1-430 | 243/528 | 0.00/0.05/0.05 | Capsid cluster; pro-, A-, and B-capsids |
| 20220615-CS02_VZV51G2_lamella2_tilt11 (16.846 Å/px) | 100-410 | 253/646 | 0.00/0.05/0.05 | Nuclear egress; C-capsid |
| 20220509_CS02_VZV51G1_lamella2_tilt03 (16.846Å/px) | 100-340 | 155/676 | 0.00/0.05/0.10 | Secondary envelopment. Glycoproteins visible; C-capsid |
| 20220507_CS02_VZV52G2_lamella3_tilt03 (16.846 Å/px) | 60-300 | 143/450 | 0.00/0.02/0.05 | VZV particle morphogenesis |
| 191017_VZV6_G4_11117b_Tilt28[E] | | 163/475 | 0.00/0.07/0.03 | Extracellular VZV particles |
| (3.5Å/px) | 1-220 | 64/475 | 0.00/0.07/0.03 | |

[A] Tilt series and tomograms are available from EMPIAR-12464.
[B] Movie slices
[C] Images captured at 1000x1000 in Zap at 0.3333; scale bar 125nm
[D] Images captured at ~1000x1000 in Slicer at 1.00; the three values represent Filtering in Fourier space/Low-frequency sigma/High-frequency cutoff
[E] Cryo-ET only, not FIB milled.

**Supplemental Table 3. Frequency of VZV capsids and their location within the infected cell.**

| Capsid Type[A] | Frequency of VZV Capsids | | | | |
|---|---|---|---|---|---|
| | Nucleus | Cytoplasm | Particle[B] | Extracellular[C] | N= |
| Procapsid | 34 | 0 | 0 | 0 | 34 |
| A | 204 | 8 | 2 | 0 | 214 |
| B | 556 | 12 | 7 | 0 | 575 |
| C | 186 | 22 | 50 | 3 | 261 |
| All | 980 | 42 | 59 | 3 | 1084 |

[A] Based on classically defined capsid morphology.
[B] Complete VZV particle in the cytoplasm found within vesicular structures.
[C] Seen in the extracellular space between VZV infected cells.

**Supplemental Table 4. The structures and proteins that comprise herpesvirus capsids.**

| Structure | Copies per Capsid | Protein | Copies per Capsid | Herpesvirus Genes [Protein[A]] | |
|---|---|---|---|---|---|
| | | | | VZV | HSV-1 |
| Penton | 11 | MCP | 55 | ORF40 | UL19 [VP5] |
| Hexon | 150 | MCP | 900 | | |
| | | SCP | 900 | ORF23 | UL35 [VP26] |
| Triplex | 320 | TRX1 | 320 | ORF20 | UL38 [VP19C] |
| | | TRX2[B] | 640 | ORF41 | UL18 [VP23] |
| CVSC | 60 | CVC1 | 60 | ORF43 | UL17 [pUL17] |
| | | CVC2[B] | 120 | ORF34 | UL25 [pUL25] |
| | | VP1/2[B,C] | 120 | ORF22 | UL36 [pUL36] |
| Portal | 1 | Portal[D] | 12 | ORF54 | UL6 [pUL6] |

[A] The associated protein names are given for HSV-1.
[B] Dimer.
[C] Only the carboxy terminal helix is found in the CVSC.
[D] Dodecamer.

239 **Supplementary Table 5. Reagents and resources.**

| REAGENT or RESOURCE | SOURCE | IDENTIFIER |
|---|---|---|
| **Virus Strain** | | |
| pOka-TK-RFP | [2] | |
| pOka-TK-GFP | [3] | |
| **Chemicals, Peptides, and Recombinant Proteins** | | |
| Minimal essential medium | Corning cellgro | 10-010-CV |
| Fetal bovine serum | Gibco | 26140-079 |
| Penicillin /Streptomycin | Gibco | 15140-122 |
| Amphotericin B | Corning cellgro | 30-003-CF |
| Nonessential amino acids | Corning cellgro | 25-025-CI |
| Gibco™ TrypLE™ Express Enzyme | Fisher Scientific | 12-605-036 |
| PLPP gel | Alveole | |
| PBS | Fisher Scientific | 21040CV |
| Deionized water | | |
| HEPES | Sigma | H3375 |
| Ethanol 100% | Gold Shield | 412804 |
| Poly-L-lysine 0.01% | Sigma | P4707 |
| mPEG-Succinimidyl Valerate, MW 5,000 Laysan Bio Inc. | Fisher Scientific | NC0107576 |
| Gelatin Oregon green 488 | Invitrogen | G13186 |
| **Deposited Data** | | |
| Original movie files, tilt series, and tomograms. | EMPIAR | EMPIAR-12464 |
| Vertex cryo-ET map; 8.3Å (EMAN2) | EMDB | EMD-49591 |
| Vertex molecular model | PDB | 9NO1 |
| 3-fold axis cryo-ET map; 8.4Å (EMAN2) | EMDB | EMD-49886 |
| Capsid cryo-ET map – 471 capsids; 12.1Å (Relion) | EMDB | EMD-49465 |
| Vertex cryo-ET map – 5,318 vertices; 10.3Å (Relion) | EMDB | EMD-49466 |
| Capsid with portal cryo-ET map – 109 capsids; 24.5Å (Relion) | EMDB | EMD-49467 |
| Portal vertex cryo-ET map – 109 vertices; 23.6Å (Relion) | EMDB | EMD-49468 |
| Portal cryo-ET map – 109 portals; 22.9Å (Relion) | EMDB | EMD-49469 |
| CAI capsid cryo-ET map – 62 capsids; 32.6Å (Relion) | EMDB | EMD-49470 |
| CAI portal vertex cryo-ET map – 62 vertices; 23.8Å (Relion) | EMDB | EMD-49471 |
| C-capsid cryo-ET map – 47 capsids; 33.6Å (Relion) | EMDB | EMD-49472 |
| C-capsid portal vertex cryo-ET map – 47 vertices; 24.3Å (Relion) | EMDB | EMD-49473 |

| | | |
|---|---|---|
| **Experimental Models: Cell Lines** | | |
| MeWo | ATCC | HTB-65 |
| **Software and Algorithms** | | |
| SerialEM v3.7 | [4] | http://bio3d.colorado.edu/SerialEM/ |
| WARP | [5] | https://warpem.github.io/warp/ |
| EMAN2 | | https://blake.bcm.edu/emanwiki/EMAN2 |
| IMOD | [6] | https://bio3d.colorado.edu/imod/ |
| 3dmod | [7] | https://bio3d.colorado.edu/imod/doc/3dmodguide.html |
| Relion v4.0 | [8,9] | https://relion.readthedocs.io/en/release-4.0/ |
| UCSF ChimeraX v1.8 | [10] | https://www.cgl.ucsf.edu/chimerax/ |
| ISOLDE | [11] | https://tristanic.github.io/isolde/ |
| VMD | [12] | https://www.ks.uiuc.edu/Research/vmd/ |
| NAMD | [13] | https://www.ks.uiuc.edu/Research/namd/ |
| Dynamo | [14] | https://www.dynamo-em.org/w/index.php |
| Dragonfly v2022.2 | Comet Technologies | https://www.theobjects.com/dragonfly |
| Illustrator CS6 | Adobe | |
| Photoshop CS6 | Adobe | |
| **Other** | | |
| Quantifoil® R 2/2, Au 300 mesh $SiO_2$ grids | Quantifoil | |
| EM GP2 | Leica | |
| Aquilos 2 cryo-FIM/SEM | ThermoFisher | |
| Talos Arctica 200kV | ThermoFisher | |
| Titan Krios 300kV | FEI | |
| Silicone sheeting 0.005” gloss | Specialty Manufacturing, Inc. | |

| | | |
|---|---|---|
| Laboratory wrapping film | Parafilm | Bemis™ PM999 |
| 10 cm plastic petri dish | Corning | 430591 |
| Negative pressure tweezers | Dumont | 0203-N3-PO |
| Glass coverslips #1.5, 24x60mm | Ted Pella, Inc. | 260423 |
| Glass-bottom dishes, 35mm, #1.5 | Cellvis | D35-20-1.5-N |
| Glass-bottom dishes, 35mm, #1.5 | MatTech | P35G-1.5-14-C |
| Plasma Etch PE-50 | Plasma Etch | PE-50 |
| Silicone stencils 4mm diameter wells | | |

#### References.

1. Sun, J., Liu, C., Peng, R., Zhang, F.K., Tong, Z., Liu, S., Shi, Y., Zhao, Z., Zeng, W.B., Gao, G.F., et al. (2020). Cryo-EM structure of the varicella-zoster virus A-capsid. Nat Commun *11*, 4795. 10.1038/s41467-020-18537-y.
2. Oliver, S.L., Yang, E., and Arvin, A.M. (2017). Dysregulated Glycoprotein B-Mediated Cell-Cell Fusion Disrupts Varicella-Zoster Virus and Host Gene Transcription during Infection. J Virol *91*. 10.1128/JVI.01613-16.
3. Yang, E., Arvin, A.M., and Oliver, S.L. (2014). The cytoplasmic domain of varicella-zoster virus glycoprotein H regulates syncytia formation and skin pathogenesis. PLoS Pathog *10*, e1004173. 10.1371/journal.ppat.1004173.
4. Mastronarde, D.N. (2005). Automated electron microscope tomography using robust prediction of specimen movements. J Struct Biol *152*, 36-51. 10.1016/j.jsb.2005.07.007.
5. Tegunov, D., and Cramer, P. (2019). Real-time cryo-electron microscopy data preprocessing with Warp. Nat Methods *16*, 1146-1152. 10.1038/s41592-019-0580-y.
6. Mastronarde, D.N. (1997). Dual-axis tomography: an approach with alignment methods that preserve resolution. J Struct Biol *120*, 343-352. 10.1006/jsbi.1997.3919.
7. Kremer, J.R., Mastronarde, D.N., and McIntosh, J.R. (1996). Computer visualization of three-dimensional image data using IMOD. J Struct Biol *116*, 71-76. 10.1006/jsbi.1996.0013.
8. Zivanov, J., Nakane, T., Forsberg, B.O., Kimanius, D., Hagen, W.J., Lindahl, E., and Scheres, S.H. (2018). New tools for automated high-resolution cryo-EM structure determination in RELION-3. Elife *7*. 10.7554/eLife.42166.
9. Scheres, S.H. (2012). RELION: implementation of a Bayesian approach to cryo-EM structure determination. J Struct Biol *180*, 519-530. 10.1016/j.jsb.2012.09.006.
10. Pettersen, E.F., Goddard, T.D., Huang, C.C., Meng, E.C., Couch, G.S., Croll, T.I., Morris, J.H., and Ferrin, T.E. (2021). UCSF ChimeraX: Structure visualization for researchers, educators, and developers. Protein Sci *30*, 70-82. 10.1002/pro.3943.
11. Croll, T.I. (2018). ISOLDE: a physically realistic environment for model building into low-resolution electron-density maps. Acta Crystallogr D Struct Biol *74*, 519-530. 10.1107/S2059798318002425.
12. Humphrey, W., Dalke, A., and Schulten, K. (1996). VMD: visual molecular dynamics. J Mol Graph *14*, 33-38, 27-38. 10.1016/0263-7855(96)00018-5.
13. Phillips, J.C., Hardy, D.J., Maia, J.D.C., Stone, J.E., Ribeiro, J.V., Bernardi, R.C., Buch, R., Fiorin, G., Henin, J., Jiang, W., et al. (2020). Scalable molecular dynamics on CPU and GPU architectures with NAMD. J Chem Phys *153*, 044130. 10.1063/5.0014475.
14. Castano-Diez, D., Kudryashev, M., Arheit, M., and Stahlberg, H. (2012). Dynamo: a flexible, user-friendly development tool for subtomogram averaging of cryo-EM data in high-performance computing environments. J Struct Biol *178*, 139-151. 10.1016/j.jsb.2011.12.017.
